# Anterior Cingulate Cortex Signals the Need to Control Intrusive Thoughts During Motivated Forgetting

**DOI:** 10.1101/2021.08.10.455844

**Authors:** Maité Crespo García, Yulin Wang, Mojun Jiang, Michael C. Anderson, Xu Lei

## Abstract

How do people limit awareness of unwanted memories? Evidence suggests that when unwelcome memories intrude, a retrieval stopping process engages the right dorsolateral prefrontal cortex (rDLPFC; Anderson et al., 2004) to inhibit hippocampal activity (Benoit and Anderson, 2012; Benoit et al., 2015; Gagnepain et al., 2017) and disrupt retrieval. It remains unknown how and when the need to engage prefrontal control is detected, and whether control operates proactively to prevent an unwelcome memory from being retrieved, or must respond reactively, to counteract its intrusion. We hypothesized that dorsal anterior cingulate cortex (dACC) achieves this function by detecting signals indicating that an unwanted trace is emerging in awareness, and transmitting the need for inhibitory control to right DLPFC (Alexander and Brown, 2011; Botvinick et al., 2001). During a memory suppression task, we measured trial-by-trial variations in dACC’s theta power and N2 amplitude, two electroencephalographic (EEG) markers of the need for enhanced control (Cavanagh and Frank, 2014). With simultaneous EEG-fMRI recordings, we tracked dynamic interactions between the dACC, rDLPFC and hippocampus during suppression. EEG analyses revealed a clear role of dACC in detecting the need for memory control, and in upregulating prefrontal inhibition. Importantly, we identified dACC contributions before episodic retrieval could have occurred (500 ms) and afterwards, indicating distinct proactive and reactive control signalling. Stronger proactive control by the dACC led to reduced hippocampal activity and diminished overall blood-oxygen-level-dependent (BOLD) signal in dACC and rDLPFC, suggesting that pre-empting retrieval early reduced overall control demands. However, when dACC activity followed the likely onset of recollection, retrieval was cancelled reactively: effective connectivity analyses revealed robust communication from dACC to rDLPFC and from rDLPFC to hippocampus, tied to successful forgetting. Together, our findings support a model in which dACC detects the emergence of unwanted content, triggering top-down inhibitory control, and in which rDLPFC countermands intruding thoughts that penetrate awareness.

## Introduction

When people suppress unwanted memories, the dACC is among the regions more active, but its contribution to inhibitory control over memory remains undefined. In non-memory contexts, major theoretical accounts agree that dACC monitors ongoing processing and detects information indicating a need to intensify cognitive control, and that dACC communicates this demand to prefrontal regions that implement control (Alexander and Brown, 2011, 2015; Botvinick, 2007; Botvinick et al., 2001; Cavanagh and Frank, 2014; Vassena et al., 2020). The conflict monitoring theory (Botvinick et al., 2001) proposes that dACC is sensitive to cognitive conflict, and that processed conflict signals initiate strategic adjustments in cognitive control to prevent future conflict. Accounts derived from the predicted response outcome model (PRO, Alexander and Brown, 2011) point out that surprising events typically increase the activation of this region, so they maintain that dACC plays a specific role in calculating surprise (Vassena et al., 2020). Following these ideas, we hypothesized that, during motivated forgetting, dACC dynamically regulates mnemonic inhibition by computing signals that indicate a need to control unwelcome content. On one hand, these warning signals may originate from reminders that foreshadow an unwanted memory’s intrusion, triggering proactive control to prevent retrieval; on the other hand, they may derive directly from an unwanted memory’s reactivation, which may elicit cognitive conflict and a need to purge the intruding memory from mind (Levy and Anderson, 2012). Specifically, when proactive control fails to prevent retrieval, intrusion-related activity would drive stronger signals in dACC as the demands for cognitive control increase, and this would initiate a reactive mechanism engaging rDLPFC and downregulating the hippocampus (Benoit et al., 2015; Gagnepain et al., 2017; Levy and Anderson, 2012). We hypothesized that dACC would transmit these signals to prefrontal regions to amplify top-down inhibition over regions driving retrieval of the offending memory.

To test these hypotheses, we acquired simultaneous EEG-fMRI recordings as participants performed a memory suppression task. This multimodal approach allowed us to relate temporally precise signatures of the need for enhanced cognitive control to BOLD signals to track dynamic interactions between the dACC, rDLPFC and hippocampus during suppression (**Figure 1**). Following an EEG-informed fMRI approach, we first tested, on a trial-by-trial basis, the coupling between BOLD signals in the foregoing regions and proactive control. We indexed proactive control via measures of frontal midline theta power and N2 amplitude arising prior to the likely recollection of the intruding memory. We hypothesized that, whereas proactive control prepares the system to inhibit hippocampal retrieval, those trials with poorer proactive control would lead to increased demands for intrusion-control later in the trial, reactively triggering elevated activity in both dACC and rDLPFC. To tie these prefrontal interactions to mnemonic control, we examined how engaging dACC/rDLPFC related the suppression of hippocampal retrieval processes by testing the coupling of prefrontal activity with theta oscillatory activity from hippocampal EEG sources. We included temporally resolved EEG source analyses to investigate whether regional modulations occurred before or after likely recollection onset. Finally, we captured the dynamics of information flow during reactive control of unwanted memories by calculating Granger causality between EEG sources. These analyses allowed us to measure whether dACC transmits control signals to rDLPFC, and whether rDLPFC, in turn, intensifies top-down inhibition of the hippocampus, facilitating motivated forgetting.

**Figure 1.**
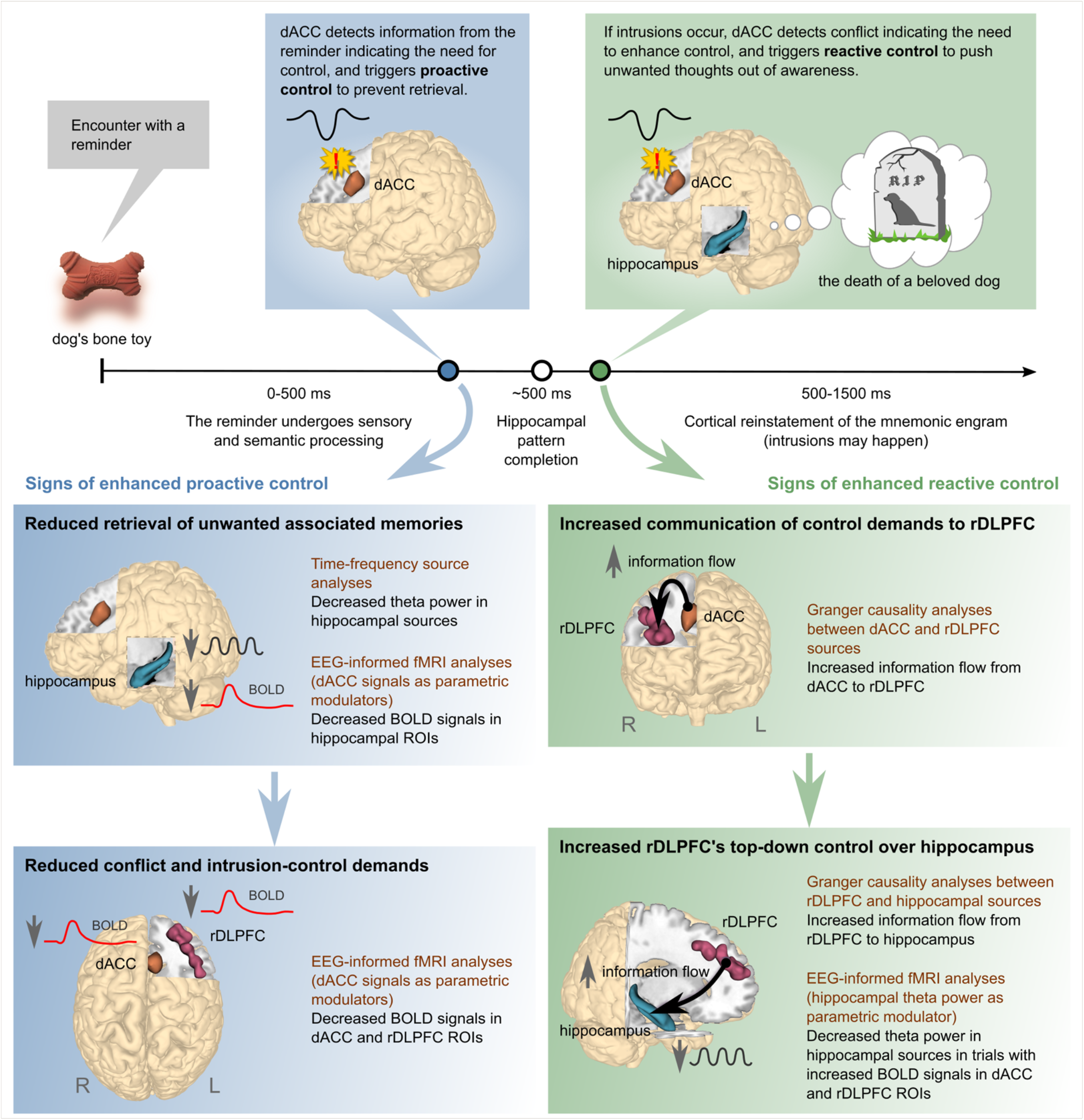
Summary of expected relationships between EEG measures and BOLD signals associated with proactive and reactive control. The upper panel represents a hypothetical timeline of brain processes after encountering a reminder (dog’s bone toy) associated with an unwanted memory (the death of a beloved dog). Proactive control (blue dot) is triggered before episodic retrieval of associated memories starts in the hippocampus, whereas reactive control (green dot) is triggered after, because of conflict generated by intrusions. The lower panels summarize the expected effects of enhanced proactive (blue boxes) and reactive (green boxes) control, according to the model. These effects imply specific relationships between EEG measures and BOLD signals, which were tested using the methods listed (brown text). Please, refer to the main text for more details.

## Results

The memory suppression task was a version of the Think/No-Think (TNT) paradigm (Anderson and Green, 2001) as shown in **Figure S1**. First, participants (n=24) encoded unrelated cue-associate word pairs and were trained to recall the associate given the cue. Then, participants entered the TNT phase, wherein they were presented with cues from studied items as reminders and directed to control the retrieval process, while we acquired simultaneous EEG-fMRI recordings. On each trial of the Think condition, participants received the cue word within a green frame and were asked to recall and think about its associate; on No-Think trials, by contrast, participants received the cue within a red frame and were asked to prevent the associate from entering consciousness. In a final phase, we performed two memory tests. On the same-probe (SP) test, participants received each cue word again and tried to recall its associate. On the independent-probe (IP) test, participants instead received a novel category name and were asked to recall a word that belonged to that category from among the studied associates. We also tested memory for items that participants learned during the training phase, but that had not appeared during the TNT phase, providing a baseline estimate of retention for items that had neither been retrieved nor suppressed.

### Behaviour and confirmatory fMRI data analyses

We replicated key findings from previous studies (Anderson and Green, 2001; Anderson and Hulbert, 2021). As expected, participants recalled fewer associate words in the No-Think than in the Baseline condition (SP test: t(23)=4.65; p*<*0.001; IP test: t(23)=4.74; p*<*0.001 and overall memory test: t(23)=6.16; p*<*0.001; **Figure S2**). The below-baseline recall performance for No-Think items reflects suppression-induced forgetting (SIF) and confirms that participants successfully engaged inhibitory control mechanisms during retrieval suppression, which impaired memory. Although memory performance was lower on the IP test than on the SP test (Test type effect: F(1,23)=94.52; p*<*0.001), SIF generalized across both tests (Condition main effect: F(1,23)=24.00; p*<*0.001; Condition*Group interaction: F*<*1)(Anderson and Green, 2001; Anderson et al., 2004). In contrast, voluntary retrieval did not affect recall of Think items compared to baseline (overall memory test: p=0.30; SP test: p=0.15) (cf. Anderson and Green, 2001). Nevertheless, using different cues at recall than those studied and practiced was detrimental for retrieval (IP test: t(23)=-2.55; p*<*0.05), as has been previously reported, which is consistent with the encoding specificity principle (for a detailed discussion, see Paz-Alonso et al., 2009).

To confirm that suppressing associate words engaged dACC and rDLPFC, we analysed fMRI data comparing the activation between NT and T trials from the TNT phase. Using a priori dACC and rDLPFC ROIs taken from a meta-analysis of 16 retrieval suppression studies (Apšvalka et al., 2020), we observed greater activity during retrieval suppression than voluntary retrieval (dACC: +6, +23, +41; p(FWE)*<*0.05, small volume corrected [SVC]; rDLPFC: +36, +38, +32; p(FWE)*<*0.01, SVC). With the opposite contrast (NT*<*T), we also confirmed decreased activity during retrieval suppression relative to voluntary retrieval in the hippocampus (left hippocampus: -33, -34, -10; p(FWE)*<*0.001, SVC; right hippocampus: +24, -25, -13; p(FWE)*<*0.05, SVC). Importantly, these deactivations were below the level observed in a perceptual baseline condition in which participants viewed unpaired single words presented within a grey frame (left hippocampus: -33, -31, -10; p(FWE)*<*0.005, SVC; right hippocampus: +30, -25, -19; p(FWE)*<*0.05, SVC), consistent with the view that retrieval suppression downregulates hippocampal activity (Depue et al., 2007; Gagnepain et al., 2017). In addition, an exploratory analysis using the overall contrast between No-Think and Think trials revealed BOLD activation patterns consistent with previous observations (for reviews, see Anderson et al., 2016; Anderson and Hanslmayr, 2014). Additional activations arose in mostly right-lateralized regions, including supplementary motor area, premotor cortex, inferior frontal gyrus, and parietal lobe, whereas additional deactivations occurred in brain areas that support the representation of visual memories, among other regions (**Table S1**; **Figure S3**).

### Early engagement of dACC’s theta control mechanism predicts reduced demands on dACC and rDLPFC for intrusion control

People who see a reminder to an unwanted thought and engage inhibitory control early enough may prevent the unwelcome memory from intruding. Achieving this form of proactive control requires a mechanism that detects the need for increased control upon seeing a reminder. In natural settings, people may learn to identify warning features of stimuli that foreshadow unpleasant thoughts and use these features to initiate proactive suppression. We hypothesized that one of the key roles of dACC during motivated forgetting is to trigger this proactive mechanism to entirely prevent awareness of unwelcome content. This proactive mechanism may be initiated in the TNT task when participants process the red No-Think cues as task signals to stop retrieval. Related to this possibility, a previous EEG study gave participants advanced warning about whether each upcoming trial required suppression or retrieval; they found that anticipating the need for retrieval suppression increased theta power in dACC and left DLPFC sources in No-Think relative to Think trials within 500 ms after the suppression task warning (Waldhauser et al., 2015). Indeed, in non-memory tasks, increased midline and prefrontal theta activity typically reflects enhanced cognitive control, and is a common mechanism by which anterior cingulate and medial prefrontal cortices detect the need for control and communicate it to lateral prefrontal cortex (Cavanagh and Frank, 2014).

Guided by these findings, we used theta power localized to dACC sources as an index of dACC engagement in upregulating inhibitory control during memory suppression and sought to relate this effect to BOLD signal in the dACC and rDLPFC. To focus on proactive control, we measured dACC-theta within an early time window after cue onset and before the likely retrieval of the associates (∼ 500 ms, Staresina and Wimber, 2019). Our hypothesis implies two main predictions about how early dACC-theta power should relate to BOLD signals. First, if dACC is engaged in proactive control, we should expect elevated theta power during the early time window. However, provided that proactive control prepares the brain for mnemonic inhibition, facilitating retrieval stopping, successful engagement of this mechanism should prevent intrusions, reducing aggregate demands on inhibitory control over the full 3-second duration of the trial (think of the adage: *“a stitch in time, saves nine”*). Therefore, although theta activity is generated by the dACC, a robust early theta response should, paradoxically, predict less aggregate dACC BOLD signal during the trial, reflecting the diminished need for intrusion control. Similarly, a robust early dACC-theta response should lead to less rDLPFC BOLD signal over the duration of the trial.

To test whether these salutary effects of proactive control emerge, we measured trial-by-trial variations of EEG theta power in dACC sources within an early time window (300-450 ms; 4-8 Hz) where frontal midline theta power started to increase in No-Think relative to Think (**Figure 2A**). We then used this measure as a parametric modulator for an EEG-informed fMRI analysis. Consistent with our first prediction, trials with enhanced early dACC-theta power were associated with reduced BOLD signal in the dACC ROI, an effect specific to memory suppression (No Think: p=0.03; No Think*<*Think: p*<*0.05; **Figure 2D**; see also **Table S2**). To test the second prediction, we used the same parametric modulator but restricted the contrasts to the rDLPFC ROI. BOLD signal in the rDLPFC ROI was also reduced in trials with enhanced early dACC theta power, and this effect was also specific to suppression (No Think: p*<*0.01; No Think*<*Think: p*<*0.05; **Figure 2D**; see also **Table S2**).

**Figure 2.**
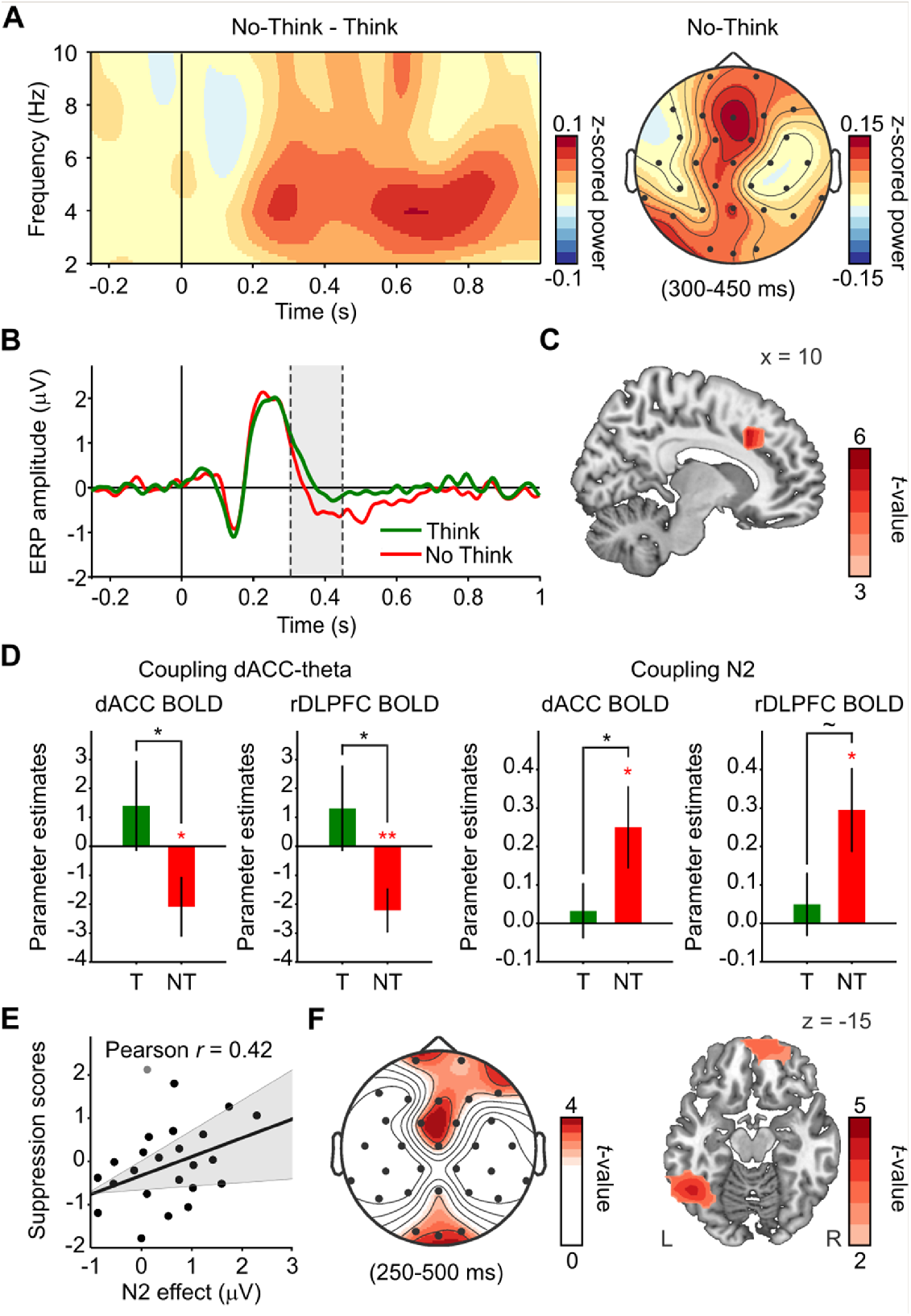
Early frontal midline theta signals in dACC during retrieval suppression reflect proactive control. **(A)** Time-frequency and scalp topographic maps showing increases in frontal midline theta power during No-Think relative to Think. There was a cluster of frontocentral channels showing significant effects between 4-6 Hz and 250-800 ms (p<0.05). We extracted theta power from dACC sources within 300-450 ms, as a measure of proactive control. **(B)** Decreased amplitude of ERP waveform in No-Think (red) relative to Think (green; p-cluster<0.05). Gray shadow indicates the time window 300-450 ms considered for N2 wave analyses. **(C)** Increased power in dACC sources during the suppression-N2 (300-450 ms; p<0.01, corrected with permutations and maximal statistic). **(D)** BOLD signals in dACC and rDLPFC were reduced in trials with increased dACC-theta power during retrieval suppression but not during retrieval. Similarly, BOLD signals in dACC and rDLPFC were reduced in trials with larger N2 signals during retrieval suppression. *p<0.05; **p<0.005; ∼ p=0.052. Error bars represent standard error of the mean. T=Think; NT=No-Think. **(E)** Individuals with larger the N2 showed more suppression-induced forgetting. Scatter plot shows a correlation between the N2 effect (ERP amplitude in Think minus No Think) and forgetting scores (z-SIF). Each participant’s forgetting z-SIF was obtained by z-scoring its suppression-induced forgetting (SIF) index relative to the SIF of all participants receiving the same items in the same conditions (counterbalancing group). This transformation isolates suppression effects, by correcting for irrelevant variability in forgetting due to differences in memorability of items across counterbalancing groups. **(F)** Scalp topo-graphic and source maps showing increases in alpha power over left occipito-temporal regions (left fusiform gyrus) between 250-500 ms in trials with large N2 signals (p-cluster<0.05).

We sought converging evidence for the role of proactive control in reducing demands on dACC and rDLPFC by focusing on a well-established ERP component related to memory suppression. Previous studies have demonstrated the suppression N2 effect (more negative-going wave in No-Think than in Think), which appears to index the engagement of early inhibitory control during suppression; it correlates with the N2 effect of motor-stopping (Mecklinger et al., 2009), with SIF, and with the consequent reduction in distressing intrusions of lab-analogue traumatic memories (Streb et al., 2016). Taking into account its early latency and medial frontal topography, we hypothesized that this suppression N2 may be partly generated in dACC and reflect an aspect of the same frontal midline theta mechanism that processes the need for control (Cavanagh and Frank, 2014) after seeing the No-Think cues. Replicating prior studies (Bergströ m et al., 2009; Chen et al., 2012; Mecklinger et al., 2009; Streb et al., 2016; Waldhauser et al., 2012), mean ERP amplitudes during No-Think trials were significantly more negative than they were during Think trials between 300-450 ms (300-400 ms: t-mean(23)=-2.20, t-cluster=-15.5, p-cluster=0.04; 350-450 ms: t-mean(23)=-2.70, t-cluster=-27.0, p-cluster=0.004) at frontocentral, right frontal and right temporal electrodes; **Figure 2B**). The N2 effect was localized at the right supplementary motor area (SMA, BA6: 10, 0, 60, p*<*0.001; **Figure 2C**), and supporting our hypothesis, at other frontal midline regions including the dACC (peak voxel: 10, 20, 40, p*<*0.001). To be consistent with how N2 is measured in other studies, we created a pooled frontocentral channel (Fz, FC1 and FC 2) and compared again the amplitudes of No-Think and Think trials within the 300-450 ms window. This frontocentral channel showed peak differences between 332-348 ms (all t(23)*<*-2.70, p-corrected*<*0.05), confirming that the N2 effect likely was generated before participants recollected the associate and agreeing with a proactive control role. Importantly, participants with a larger N2 effect (more negative going N2 in No-Think relative to Think trials) showed higher SIF scores (r=0.42; p*<*0.05; 1 outlier; **Figure 2E**); indeed, only high forgetters showed differences in amplitude during this time window (332-348 ms, all t(11)*<*-2.88, p-corrected*<*0.05; **Figure S4**) but not low forgetters (all t(11)*>*-1.66, p-corrected*>*0.24). Our results confirm previous findings and suggest that the N2 effect reflects early control processes in dACC that facilitate memory inhibition (Bergströ m et al., 2009; Mecklinger et al., 2009).

If the N2 reflects early inhibitory control, increases in this component, just as with dACC-theta power, should be negatively related to BOLD signal in both the dACC and rDLPFC. To test this, we extracted trial-by-trial variations of the N2 amplitude at the frontocentral channel. Then, we used this measure to build a parametric modulator for an EEG-informed fMRI analysis. Consistent with our hypothesis, trials with larger (more negative) frontocentral N2 amplitudes were accompanied by reduced BOLD signal in the dACC ROI (p*<*0.05) specifically in No-Think trials (p*<*0.05; **Figure 2D**; see also **Table S3**). Similarly, trials with larger (more negative) frontocentral N2 amplitudes during memory suppression were associated with reduced BOLD signal in the rDLPFC ROI (p*<*0.05), although the differences relative to voluntary retrieval did not achieve significance (p=0.052). Taken together, the timing of the N2 component and the negative relationship to overall BOLD signal across the full duration of the trial suggest that the N2 effect, like dACC-theta, indexes proactive inhibitory control over memory. This proactive mechanism contributes to stopping retrieval processes, reducing the occurrence of intrusions, and pre-empting any need for further engagement of dACC and rDLPFC during the suppression trial.

To further scrutinise this hypothesis, we tested whether the N2 was associated with EEG oscillatory markers of elevated cognitive control. Firstly, we expected to link the N2 with early increases in frontal midline theta activity; and secondly, we expected to find enhanced alpha/beta band activity in regions involved or under top-down inhibitory control (Castiglione et al., 2019; Fellner et al., 2020; Waldhauser et al., 2015). To test this, we split sensors’ TFRs on No-Think trials based on the amplitude of the frontocentral N2 as an index of proactive control. Trials with larger (more negative) N2 amplitudes showed, in addition to frontocentral theta power increases, increased alpha power relative to trials with smaller N2 amplitudes. Effects were maximal at frontal (4-8 Hz: 200-500 ms; 9-12 Hz, 350-450 ms; all t*>*2.46, t-cluster=226.9; p-cluster=0.01) and occipitoparietal sensors (9-12 Hz: 250-500 ms; all t*>*2.55, t-cluster=132.8; p-cluster=0.057; **Figure 2F**). Alpha power differences were maximal in two clusters of brain sources, one including left temporal and occipital areas (peak voxel: -50, -50, 0, BA37; t=4.54, p-cluster=0.038) and another one maximal in right medial superior frontal gyrus (peak voxel: 10, 70, 10, BA10; t=4.07, p-cluster=0.019). These findings suggest a possible mechanistic link between frontal midline theta and posterior alpha power increases (Waldhauser et al., 2015). This suggests that successfully pre-empting intrusion-related conflict via early proactive control might be partly achieved by increasing local inhibition in visual cortical areas and suppressing cortical memory representations (Gagnepain et al., 2014).

### Delayed evoked responses in dACC during suppression may reflect the detection of control demands caused by intrusions

When early control fails to suppress cue-driven retrieval, unwanted memories may intrude into awareness, creating cognitive conflict, driving higher demands on inhibitory control over memory (Levy and Anderson, 2012). We hypothesized that, in addition to its role in signalling early control demands in response to the task cues, dACC detects intrusions as ‘ alarm’signals that indicate the need to further increase inhibitory control (Alexander and Brown, 2011). If intruding recollections trigger such a delayed control response, modulations in the evoked activity of dACC should arise after the latency where recollection could start (∼ 500 ms, Staresina and Wimber, 2019). To test this prediction, we extracted source time courses (0.5-30 Hz) from all voxels within our dACC ROI and compared the mean amplitudes observed during No-Think and Think trials across all time points after cue onset (0-1 s). Accordingly, dACC amplitude was more negative during No-Think trials than during Think trials between 548-708 ms (t-mean(23)=-2.51, t-cluster=-793.6, p-cluster=0.0064; **Figure 3A**). Amplitude differences were only significant in dACC sources oriented radially to the top of the head, consistent with the topography of ERPs related to novelty, conflict, or error processing (Cavanagh et al., 2012). These ERPs are thought to be part of the same frontal midline theta mechanism for realizing the need for cognitive control (Cavanagh and Frank, 2014); therefore, we expected that these delayed control signals in dACC particularly would involve theta band activity. Indeed, amplitude differences were explained by delta/theta band evoked activity (0.5-8 Hz: 428-728 ms, t-mean(23)=-2.51, t-cluster=-1511, p-cluster=0.0046) and were not significant for alpha/beta evoked activity (8-30Hz: all p-clusters*>*0.33).

**Figure 3.**
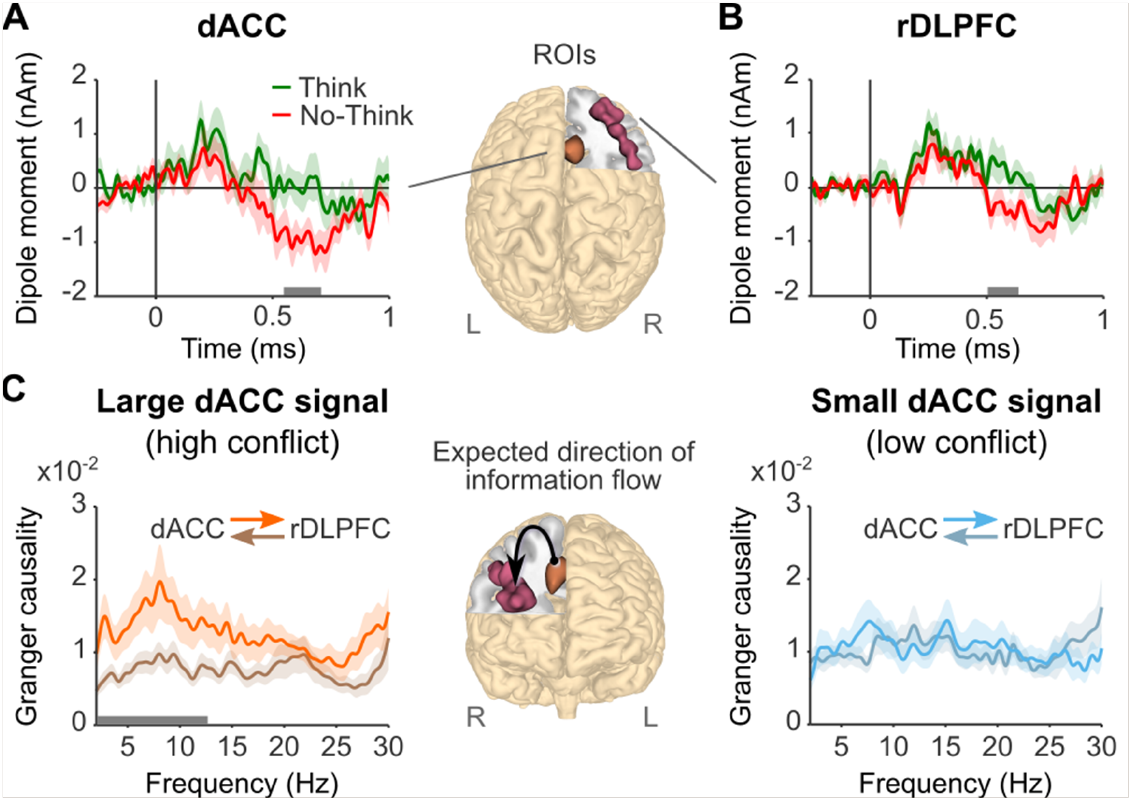
Time courses and Granger Causality analysis of delayed dACC and rDLPFC effects. **(A** and **B)** Mean evoked activity (0.5-30 Hz) in dACC **(A)** and rDLPFC **(B)** showing significant differences between each condition in a delayed time window (548-708 ms and 504-640 ms, respectively). Horizontal grey bars indicate the significant time windows (p-cluster<0.05). EEG source time courses were reconstructed from the ROIs showed in the brain representations. **(C)** Granger causality spectra of information flow between dACC and rDLPFC sources (450-1450 ms) in trials associated with high and low conflict (large or small dACC activity in the 428-728 ms window, 0.5-8 Hz). In trials with more conflict-related activity, the information flow goes primarily from dACC to DLPFC between 2-12.5 Hz. Horizontal grey bar indicates the significant frequency window (p-cluster<0.05). Light shadowed areas represent standard error of the mean.

If intruding recollections drive these delayed dACC evoked responses, these signals should be more evident for individuals likely to experience intrusions. Agreeing with this, only low forgetters, who should have experienced more overall conflict due to intrusions (Gagnepain et al., 2017), showed more negative amplitudes in the No-Think than the Think condition between 552-704 ms (t-mean(11)=-2.68, t-cluster=-769.8, p-cluster=0.0044). In contrast, high forgetters, who were more efficient at implementing inhibitory control and forgetting the associates, showed no amplitude differences in this later window. These findings are in line with our hypothesis that dACC is also involved in the late adjustment of inhibitory control when unwanted memories emerge. Moreover, they suggest that late evoked theta responses in dACC reflect cognitive conflict and control demand signals driven by intrusions.

### Intrusions trigger dACC to communicate the need to increase inhibitory control to rDLPFC through a theta mechanism

Once an intrusion is detected, the need for increased control should be transmitted to prefrontal areas that implement mnemonic inhibition. We hypothesized that dACC communicates an intrusion-control signal to rDLPFC through a mechanism of interareal neural coupling mediated by theta band activity (Cavanagh and Frank, 2014; Smith et al., 2019). To test this idea, we first investigated whether rDLPFC evoked activity showed modulations reflecting the reception of dACC signals. We reconstructed source time courses (0.5-30 Hz) from all voxels within the rDLPFC ROI and compared their mean amplitudes across No-Think and Think trials for all time points after cue onset (0-1 s). rDLPFC amplitude was more negative in No-Think than in Think trials at latencies overlapping the dACC effect associated with putative intrusion control (sources with left-right orientation: 408-588 ms, t-mean(23)=-2.56, t-cluster=1.374, p-cluster=0.008; sources with inferior-superior orientation: 504-640 ms, t-mean(23)=-2.53, t-cluster=-1.425, p-cluster=0.016; **Figure 3B**). Then, we performed a whole-brain source analysis within dACC’s delayed control window (0.5-8 Hz; 428-728 ms) to verify that these modulations were regionally specific and not caused by stronger nearby sources. We expected that, if dACC and rDLPFC were particularly engaged in control processes triggered by intrusions, these regions should show activity differences significantly larger than those shown prior to cue onset. Agreeing with this, two frontal clusters exhibited above-baseline effects: one in the right hemisphere comprising superior frontal gyrus, middle frontal gyrus and anterior cingulate cortex (peak voxel: 30, 30, 40, BA8; t=3.75, p-cluster=0.0086; **Figure S5**) and another in the left hemisphere comprising inferior frontal gyrus and insula (peak voxel: -50, 20, 0, BA47; t=3.84, p-cluster=0.037). We confirmed that dACC and rDLPFC ROIs showed significant effects (p*<*0.05, corrected with permutations and maximal statistics).

Finally, we looked for functional coupling between dACC and rDLPFC, indicating the transmission of the control demands. We investigated the directionality of the interaction between dACC and rDLPFC by applying nonparametric Granger causality analyses to a 1-s window after the N2 (450-1450 ms). In trials with larger intrusion-control signals (more negative evoked activity in dACC), the information flow in the direction of dACC to rDLPFC was higher than that from rDLPFC to dACC for theta and alpha frequency bands (2-12.5 Hz, p=0.0064, corrected with cluster-based permutation test). In trials with smaller dACC signals, there was no preferred direction in the information flow (p-cluster*>*0.44; interaction effect: p-cluster=0.09; **Figure 3C**). Our results support our hypothesis, showing that dACC and rDLPFC were maximally engaged around the time that unwanted memories may have been retrieved. Strikingly, stronger responses in dACC generated Granger causal influences on rDLPFC activity within the theta and alpha frequency bands, consistent with a process communicating the need to intensify memory control.

### Early control processes involving dACC contribute to later hippocampal downregulation

An intruding memory may not endure very long in awareness if inhibitory mechanisms already have been prepared at the time the intrusions occur. We hypothesized that proactive control early in the trial would facilitate the later down-regulation of memory-related networks, and that decreased theta oscillatory activity associated with retrieval (Waldhauser et al., 2015) should reflect the impact of this suppression. We tested this hypothesis by median-splitting No-Think trials according by their N2 amplitude (proactive control) and then looking for reflections of suppressed activity in sensor TFRs by applying the contrast larger*<*smaller N2 amplitude. Indeed, trials with larger N2 amplitudes showed reduced theta oscillatory power between 650-1850 ms after cue onset in left frontal, central and parietal sensors, relative to those with smaller N2 amplitudes (**Figure 4A**; 4-9 Hz; all t*<*-2.42, t-cluster=-493.9; p-cluster*<*0.001). In trials with smaller N2 amplitudes (where proactive control was reduced) theta oscillatory activity persisted above baseline levels for the whole epoch. Consistent with our hypothesis, the greatest reduction in theta was localized in the left hippocampus (peak voxel: -30, -30, -10; t=-4.56, p-cluster=0.0038; **Figure 4B**), but both hippocampi showed theta power decreases in ROI TFR analyses (left hippocampus: all t*<*-1.97; p-cluster*<*0.005; right hippocampus: all t*<*-1.84, p-cluster=0.014; **Figure 4C**).

**Figure 4.**
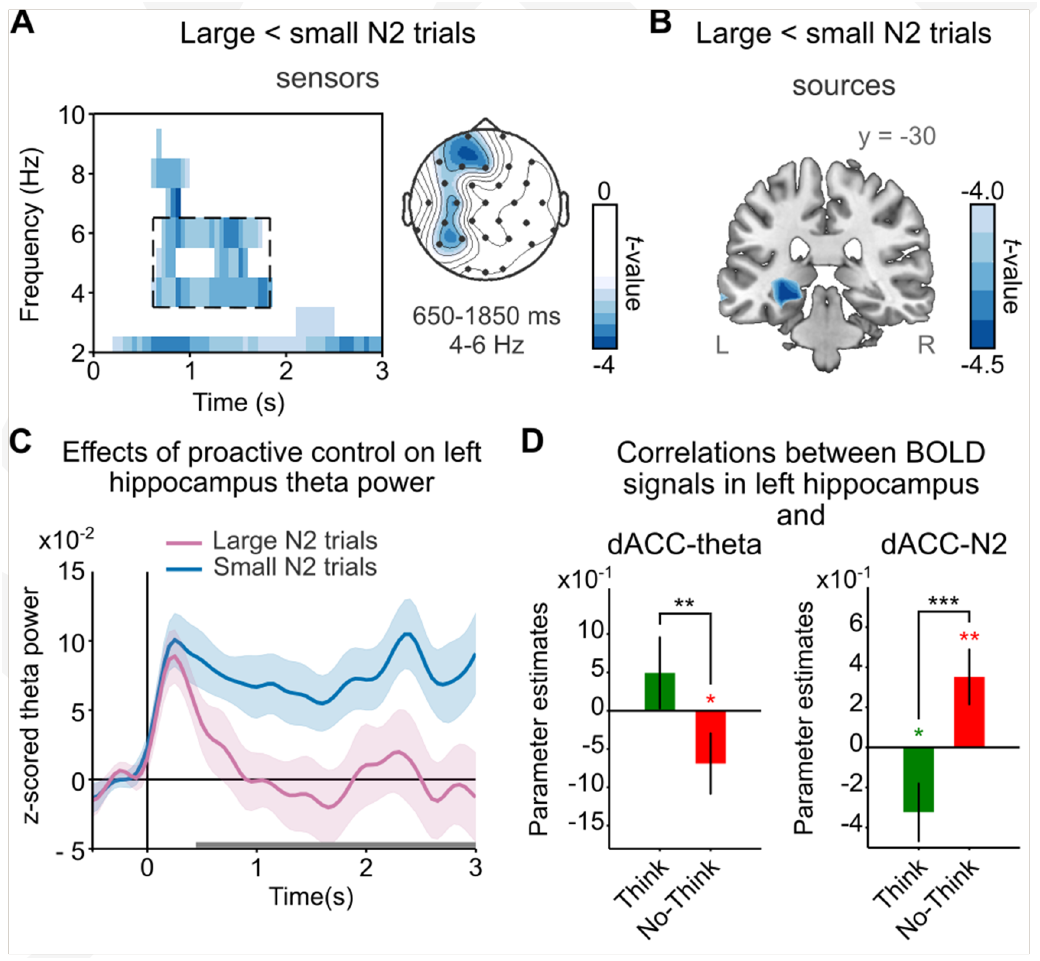
Effect of proactive control on theta power activity and BOLD signals in the hippocampus. **(A)** TFRs from all sensors showing decreased theta power in trials associated with strong proactive control (large N2 signal) relative to trials with weak proactive control (small N2 signal; p-cluster<0.05). Right panel shows the scalp topography of the effects in the framed time window (4-6 Hz, 650-1850 ms). **(B)** Sources of the peak theta power decreases were localized in left hippocampus (p-cluster<0.05). **(C)** ROI average of hippocampus z-scored power time courses for trials associated with strong and weak proactive control. Horizontal grey bars indicate the time window of significant differences between conditions in the ROI time-frequency analysis (p-cluster<0.05). **(D)** BOLD signals in the hippocampus were reduced in trials with increased dACC theta power (300-450 ms) during retrieval suppression but not during retrieval. Similarly, BOLD signals in the hippocampus were reduced in trials with larger (more negative) dACC N2 amplitudes during retrieval suppression but not during retrieval. *p<0.05; **p<0.005; ***p<0.0005. Light shadowed areas **(C)** and error bars **(D)** represent standard error of the mean.

These findings suggest that early proactive control processes contributed to the later stopping of hippocampal retrieval. To verify the localization of these effects to the hippocampus, we performed several additional analyses. First, we related EEG theta activity reconstructed from hippocampal sources to hippocampal BOLD signal. EEG-informed fMRI analysis using mean hippocampal theta power (4-6 Hz, 650-1850 ms) as a parametric modulator revealed a positive correlation between theta power and hippocampal ROIs BOLD signals (right: p*<*0.005, uncorrected; SVC, p[FWE]=0.067; left: p*<*0.05, uncorrected; **Figure 5A**) across all trials, consistent with the hypothesis that our hippocampal theta sources reflected hippocampal processing. Next, we sought converging evidence of proactive mnemonic control by relating trial-by-trial variations in early dACC-theta power to hippocampal BOLD signals during retrieval suppression. We found that trials with increased dACC-theta power showed stronger hippocampal deactivations, specifically during memory suppression (**Figure 4D**). We confirmed with an alternative EEG-informed fMRI analysis that more negative N2 amplitudes, extracted trial-by-trial from the dACC ROI, also correlated with reduced hippocampal BOLD signals, again specifically during memory suppression (SVC, p[FEW]*<*0.05; **Figure 4D**). Together, these parallel findings strongly support our hypothesis that dACC triggers proactive inhibitory control to stop retrieval of unwanted memories by the hippocampus.

**Figure 5.**
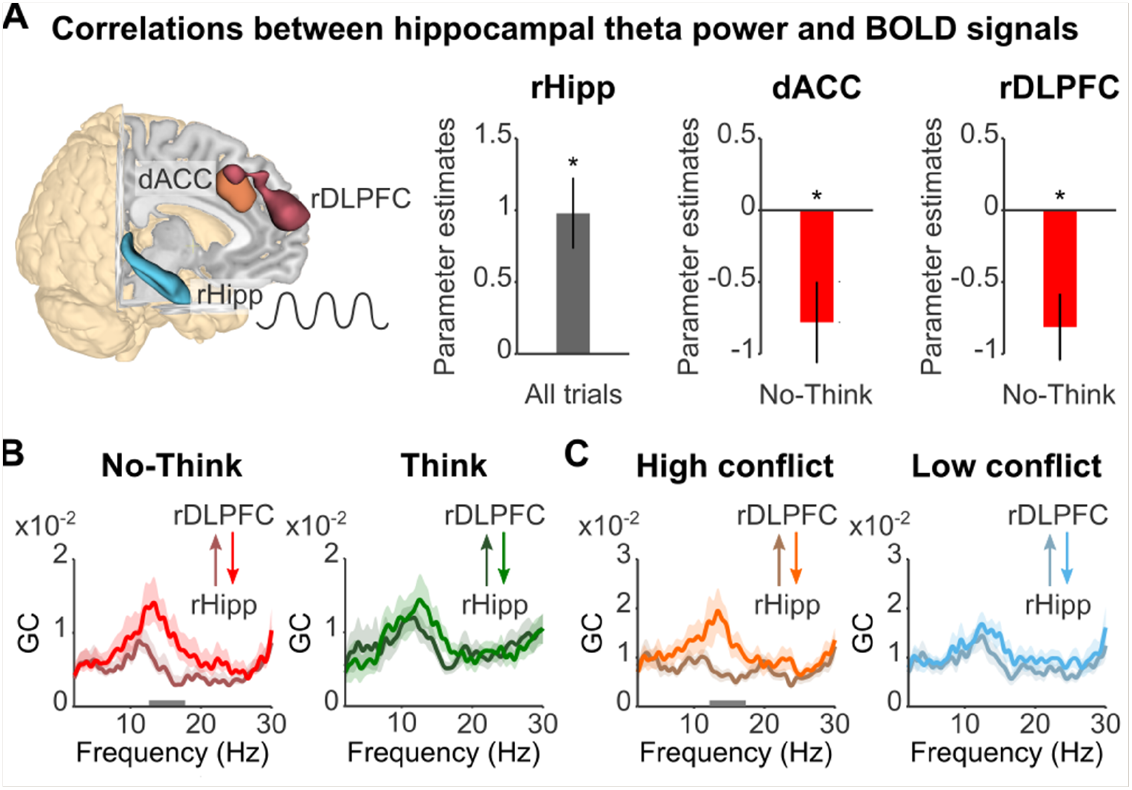
Downregulation of hippocampus by reactive control. **(A)** EEG-informed fMRI parametric analyses using hippocampal theta power as regressor. Voxels in dACC (orange) and rDLPFC (burgundy) showing significant (p<0.005) negative correlation with theta power reconstructed from sources of right hippocampus ROI (cyan). Bar plots show that theta power in the right hippocampus was (i) positively coupled to BOLD signal in this region and (ii) reduced in trials with increased dACC and rDLPFC BOLD signals during retrieval suppression. **(B** and **C)** Granger causality spectra of information flow between rDLPFC and right hippocampus sources in No-Think and Think **(B)** and trials associated with high and low conflict (large or small dACC activity in the 548-708 ms time window; **C**). In No-Think and in trials with more conflict-related activity, the information flow goes primarily from rDLPFC to right hippocampus in low beta band. Horizontal grey bars indicate the significant frequency windows (p-cluster<0.05). Error bars **(A)** and light shadowed areas **(B**,**C)** represent standard error of the mean. GC=granger causality values.

The delayed latency (650-1850ms) of the theta suppression effect suggests that although proactive control processes may help to prevent intrusions, they also enhance readiness to purge intruding thoughts from awareness when they do occur, overlapping with and likely facilitating hippocampal downregulation by reactive control processes. To examine whether a reactive hippocampal downregulation arose, we analyzed the time course of hippocampal theta activity as an index of hippocampal retrieval. Specifically, we examined those trials associated with higher putative intrusion control, to determine the presence of (a) an early increase in hippocampal retrieval activity followed by hippocampal activity suppression. To test this, we divided No-Think trials according to the power of the intrusion-control signals they exhibited in dACC (2-8 Hz; 428-728 ms). In trials with larger intrusion control signals, hippocampal theta power (2-8 Hz) was increased only during the first half of the epoch (left: 300-1600 ms, t-mean=3.16, t cluster=p-cluster*<*0.001; right: 0-1700 ms, t-mean=2.81, t cluster=p-cluster*<*0.01), relative to trials showing smaller intrusion-control signals. The fact that no significant differences emerged in hippocampal theta power during the second half of the epoch is consistent with reactive suppression of hippocampal theta in response to an intrusive recollection.

### RDLPFC down-regulates the hippocampus in response to dACC intrusion-related control signals

The foregoing findings suggest that early and delayed control mechanisms during memory suppression contribute to hippocampal down-regulation, which is expressed as theta power decreases. Particularly, when early control fails to prevent unwanted memories from intruding, dACC generates signals indicating the need for intrusion-control. These signals may dynamically adjust inhibitory control mechanisms in the rDLPFC to downregulate hippocampal activity (Benoit et al., 2015) to purge the intruding memories. Consistent with this intrusion-purging hypothesis (Levy and Anderson, 2012), during suppression, increased BOLD signals in rDLPFC and dACC were coupled to decreased theta power in bilateral hippocampal sources after the onset of recollection (650-1850ms) (DLPFC and right hippocampus: SVC, p[FWE]=0.02; and left hippocampus: p*<*0.005, uncorrected; dACC and right hippocampus: SVC, p FEW=0.01), and the latter theta suppression effect was itself associated with hippocampal downregulation (**Figure 5A**). These findings suggest that hippocampal theta power decreases (observed after retrieval onset) may arise from increased prefrontal inhibitory control.

To examine whether rDLPFC and dACC activity during intrusion-control may be causally related to hippocampal theta decreases, we used nonparametric Granger causality focusing on a 1-s window after the N2 (450-1450 ms). First, we investigated the direction of information flow between rDLPFC and hippocampus. This analysis revealed a higher top-down than bottom-up information flow during retrieval suppression in the theta band (DLPFC to left hippocampus, 2.0-6.5 Hz, p=0.021, corrected with cluster-based permutations; interaction effect: 2-5.6 Hz, p-cluster*<*0.05) and in the low beta band (DLPFC to right hippocampus, 12.6-17.8 Hz, p=0.023, corrected with cluster-based permutations; **Figure 5B**). Importantly, the increased top-down information flow in the beta range specifically arose during trials with larger intrusion-control signals in dACC (12.2-17.4 Hz, p-cluster=0.023; interaction effect: 12.3-15.4 Hz, p-cluster=0.023; **Figure 5C**), consistent with the possibility that dACC triggered the rDLPFC top-down mechanism that caused hippocampal down-regulation. To test this idea, we identified trials in which the suppressive impact of the hippocampus was clear and examined the influence of dACC on rDLPFC; we split No-Think trials according to the mean theta power in hippocampal sources (4-6Hz, 650-1850 ms) and computed Granger causality analyses within that window. We found that trials with reduced hippocampal theta showed a higher information flow in the theta band (4-9.73 Hz, p-cluster*<*0.05) from dACC to rDLPFC than in the opposite direction, consistent with a role of dACC in triggering elevated control. Together, these findings point to a late influence of the rDLPFC on the hippocampus and suggest that beta oscillations mediate a reactive top-down inhibitory control mechanism triggered in response to intrusions detected by the dACC.

Top-down inhibitory control from the rDLPFC to the hippocampus may facilitate forgetting of the suppressed memories (Anderson and Hulbert, 2021; Apšvalka et al., 2020; Benoit et al., 2015; Gagnepain et al., 2014). We tested this possibility by dividing participants into those who showed higher or lower SIF scores. High forgetters showed a greater top-down than bottom-up information flow from rDLPFC to right hippocampus for suppression items that they successfully forgot, relative to those that they remembered (19.8-23.0 Hz, p-cluster=0.031). In contrast, less successful forgetters showed the opposite pattern, with a higher top-down than bottom-up information flow for suppression items that they later remembered (11.6-17.6 Hz, p-cluster=0.006). These findings suggest that in low forgetters, the reactive engagement of top-down inhibitory control in response to intrusions is related to persisting memory for intruding thoughts.

## Discussion

Our findings reveal a central role of dACC in triggering inhibitory control that causes motivated forgetting. The data suggest that dACC can signal the need for inhibition proactively, in response to environmental cues, or reactively, to counter intrusive thoughts themselves. Two key findings support dACC’s proactive role in preventing retrieval. First, frontal-midline theta mechanisms partly originated in dACC and emerged in an early window (300-450 ms after No-Think task cue) prior to the likely onset of episodic recollection; and second, later in the trial, these early effects were associated with reduced BOLD signals and theta power in the hippocampus, consistent with reduced retrieval activity. These findings suggest that rapidly detecting the need for suppression from an environmental signal (in this case, the task cue) engages proactive control by dACC to prevent intrusions. Consistent with this interpretation, trials with increased early theta signals from dACC were accompanied by lower overall BOLD activations of dACC and rDLPFC, reflecting reduced conflict processing and lower demands on prefrontal control when recollection was quickly mitigated. Thus, rapid early control was beneficial; as they old adage says, *“a stitch in time, saves nine*.*”* These effects echo those of a prior fMRI study investigating the benefits of forgetting on neural processing during a retrieval-induced forgetting task (Kuhl et al., 2007). That study showed that, as competing memories were suppressed across retrieval practice trials, the demands to detect and overcome conflict were reduced, and so activations in ACC and lateral prefrontal cortex declined. Similar conflict reduction benefits (associations between successful memory control and reduced conflict processing) have been shown in a range of studies (see Anderson and Hulbert, 2021 for a review).

Importantly, our results suggest that proactive control did not simply prevent retrieval but also facilitated forgetting. Several observations support this conclusion. Firstly, individuals with stronger signatures of proactive control showed superior suppression-induced forgetting, complementing earlier findings linking the suppression N2 (i.e., more negative-going frontocentral ERPs around 300-450 ms Mecklinger et al., 2009) to better SIF (Bergströ m et al., 2009), and to fewer distressing intrusions after analogue trauma (Streb et al., 2016). Second, proactive control engages processes that could plausibly contribute to forgetting. For example, the N2 was partly generated in dACC and associated with early (*<*500 ms) frontal midline theta power increases, an activity pattern likely reflecting the detection and communication of the need for control to prefrontal cortex (Cavanagh and Frank, 2014). These mechanisms could trigger early inhibitory control targeting regions supporting memory. Indeed, early N2-associated increases in alpha oscillatory power (a typical correlate of cortical inhibition Klimesch, 2012) in left fusiform gyrus may be an example: given that down-regulation of this fusiform region by DLPFC during suppression is known to disrupt visual memory traces of the associates (Gagnepain et al., 2014), elevated alpha power in visual cortex may both limit the reinstatement of associated words and suppress their memory traces, resembling the downregulation of item-specific memories (Fellner et al., 2020). We also linked early proactive control signals to decreased hippocampal theta oscillatory power during a later time window (650-1850 ms), an effect previously linked to successful suppression-induced forgetting [19]. The current hippocampal modulations started during earlier time windows (∼ 300 ms), suggesting that early control reduced hippocampal theta oscillations, affecting later hippocampal-cortical theta networks. Given that hippocampal pattern completion appears to begin around 500 ms after cue onset (Staresina and Wimber, 2019), the 650-1850 ms window offers an opportunity for rDLPFC action over the hippocampus (Apšvalka et al., 2020; Benoit and Anderson, 2012; Benoit et al., 2015; Depue et al., 2007; Gagnepain et al., 2017) to inhibit memory traces, stopping intrusions. Indeed, we found significant causal influence of rDLPFC on both hippocampi within 450-1450 ms only during suppression. Together, these findings suggest that proactive control facilitates forgetting by increasing inhibition in regions where memories would be reactivated, and, by magnifying the impact of intrusion-control mechanisms (see also Hanslmayr et al., 2009; Waldhauser et al., 2015).

Critically, however, the dACC also triggered inhibitory control reactively in a time window during which unwanted memories could have been involuntarily retrieved. Our findings suggest that dACC detects conflict caused by intruding memories and communicates the need for control to rDLPFC, which in turn increases topdown inhibition over the hippocampus. Supporting this conclusion, we found, during retrieval suppression, peak dACC and rDLPFC source activities in a delayed time window, coinciding with the likely onset of conscious retrieval. dACC activity in this window (552-704 ms) was particularly large for low forgetters, suggesting that these signals reflected intrusion-related conflict. Moreover, dACC amplitude increases were due to delta/theta activity, which is a preferred rhythm for the coherent firing of dACC and DLPFC’s neurons during conflict processing, and for cross-areal coordination to implement control (Smith et al., 2019). Precisely, using dACC amplitude in this window (0.5-8 Hz; 428-728 ms) as a proxy for conflict, we found that high conflict was associated with high information flow from dACC to rDLPFC in delta/theta band and from rDLPFC to right hippocampus in beta band. The greater the activation in dACC and rDLPFC, and the stronger their theta-mediated communication, the stronger was the implementation of control over hippocampal activity (reflected by decreases in retrieval-related theta).

Prefrontal control over hippocampal activity, once triggered by dACC, appears to be achieved via beta phase synchronization. During retrieval suppression, the rDLPFC increased its communication with the right hippocampus in the low beta band, an effect not found during retrieval trials. This higher information flow, moreover, arose specifically on high conflict trials, consistent with the need to intensify top-down control to purge intrusions from awareness. Elevated communication with the hippocampus in the beta band during intrusions integrates prior evidence that had separately illustrated the importance of intrusions and beta activity in inhibitory control over memory. On the one hand, fMRI studies have found that intrusions elicit greater rDLPFC activation (Benoit et al., 2015), stronger hippocampal suppression (Benoit et al., 2015; Gagnepain et al., 2017; Levy and Anderson, 2012), and more robust fronto-hippocampal interactions to suppress retrieval (Benoit et al., 2015; Gagnepain et al., 2017), although the oscillatory mechanisms of this intrusion effects were not established. On the other hand, the importance of beta oscillations to memory inhibition has grown increasingly clear but has not linked this activity to reactive control over intrusions. For example, it has previously been shown that (a) at the scalp level, retrieval suppression increases long-range synchronization in low beta frequency band (15-19 Hz) (Waldhauser et al., 2015), suggesting that this rhythm (together with alpha) might mediate top-down control; (b) at the intracranial level, directed forgetting instructions elicit DLPFC-hippocampal interactions in the low beta range (15-18 Hz) (Oehrn et al., 2018), with greater information flow from DLPFC to hippocampus when people were instructed to forget an item, but not when instructed to remember it; and (c)stopping actions and stopping retrieval elicit a common right frontal low beta component (Castiglione et al., 2019). Taken together with the current results, these findings point to a key role of beta oscillations in the top-down control over hippocampal processing by the rDLPFC, a demand that is especially acute during the reactive control of intrusions.

Overall, the validity of our model should be further investigated with high-density EEG, magnetoencephalography, and direct electrophysiological recordings in the involved regions. EEG and fMRI recordings might display spurious correlations introduced by head movements (Fellner et al., 2016), although control analyses indicate that our effects are not due to movement (see **Supplemental Information**). EEG motion artifacts were spatially filtered out with beamforming when using source activities as parametric modulators or removed by discarding outliers. On the other hand, source reconstruction accuracy is known to be affected by low signal-to-noise ratio (especially for deep sources in the hippocampus), among other factors. We minimized this problem by following methodological recommendations (Ruzich et al., 2019).

In summary, this study provides evidence that theta mechanisms in dACC are key to triggering inhibitory control by rDLPFC during motivated forgetting. These mechanisms can be proactively engaged by external warning stimuli, helping to rapidly pre-empt unwanted thoughts. Additionally, they are strongly activated during a later time window after hippocampal retrieval likely has occurred, consistent with a reactive control response to intrusions that enhances hippocampal downregulation by the rDLPFC. This impact of prefrontal cortex on hippocampal activity is achieved by rDLPFC-hippocampal beta interactions critical to clearing the mind from unwanted thoughts and to hastening the demise of memories we would prefer not to have.

## Materials and Methods

### Participants

A total of 24 participants (12 females, mean ± SD age, 21.4 ± 2.0 years) were recruited through the Southwest University undergraduate participant pool. All had normal or corrected-to-normal vision and had no history of psychiatric or neurological illness. To check whether the participants had adequate sleep prior to the experiment, they answered questions about their sleep state upon arrival at the laboratory; all of them were in line with our requirements. None of the participants had experienced the experimental task before. The Ethics Committee of Southwest University approved the study. Written informed consents were obtained from all participants according to the declaration of Helsinki after detailed explanation of the experiment protocol. All the participants received monetary compensation after their participation.

### Stimuli

128 neutral words were selected from the Thesaurus of Modern Chinese to form 68 pairs of weakly related words. Within each pair, a word was used as cue and the other word as associate. Each associate word was a member of a unique semantic category, so it could be later recalled using that extra-list category name as cue. 48 word pairs were divided into three sets of 16 word pairs, which rotated across participants through the conditions (Think, No-Think, and Baseline). The remaining 16 pairs were used as fillers for practice. 8 additional single words were included during the TNT task in a Perceptual baseline condition.

### Experimental paradigm

The experiment consisted of three phases: study, TNT, and final memory test.

#### Study phase

First, participants studied all the 64 cue-associate word pairs. On each trial, both words were displayed visually, side by side, on a black background for 5 s. Each trial was separated by an inter-stimulus interval (ISI) with a fixation cross for 600 ms. Then, participants were trained to recall the associate words given the cues. On each trial, a cue appeared at the center of the screen for 5 s, and participants were asked to recall and say out loud the corresponding associate word. Participants’ responses were recorded. After every trial, the associate word was displayed as feedback for 2 s. All word pairs were repeatedly trained until participants correctly provided at least 50% of all the associate words. Finally, participants were tested again by displaying each cue, but the associate feedback was omitted. This test was used to identify the word pairs that participants successfully learned before entering the TNT phase and restrict (conditionalize) the analyses to those trials corresponding to learned associations.

#### TNT phase

Participants performed this part of the experiment inside the fMRI scanner and stimuli were displayed on a back-projection screen mounted above participants’ heads. At the beginning of this phase, participants practiced the task on 16 fillers. Afterwards, short diagnostic questionnaires were administered to assess whether participants understood the instructions, and questions were clarified. The proper task was divided into 6 blocks separated by 1-min breaks. Each block consisted of 80 trials, where all cues from the Think (16) and No-Think (16) conditions, together with Perceptual baseline words (8), were presented twice. In sum, each cue word was presented 12 times during this phase. Each trial started with a fixation cross (variable ISI between 500 ms and 1200 ms). Then, a cue word appeared within a coloured frame for 3 s. The trial ended with a blank screen (ISI=1.5 s). For cues within green frames (Think), participants were asked to think of the associate word and keep it in mind while the cue was on the screen. For cues within red frames (No-Think), participants were asked to pay full attention to the cue and prevent the associate word from coming into mind during the whole trial. Instructions encouraged that participants followed a direct suppression strategy (Benoit and Anderson, 2012; Bergströ m et al., 2009), by emphasizing that they should suppress retrieval and avoid replacing the associate with alternative words or thoughts. For words within a grey frame, participants were just asked to pay attention to them.

#### Final memory test

Memory for all studied word pairs was evaluated, including Baseline items that were excluded from the TNT phase. Until this phase, participants were unaware of a final memory assessment to prevent the influence of anticipatory mechanisms on forgetting scores and were initially told that the experiment was about attention and their ability to ignore distraction. Participants performed two types of final tests: same probe (SP) and independent probe (IP), with the order of these tests counterbalanced across participants. On the SP test, each cue was presented again, and participants were asked to recall and say out loud the associate word, as they did during the training phase. On the IP test, each unique category name was given as cue and participants were asked to recall and say out loud a member of this category, from those associate words they initially studied. At the end of the experiment, participants completed a questionnaire to determine whether they followed the instructions to suppress retrieval during No-Think trials.

### Equipment

#### EEG

EEG data were recorded by 32 Ag/Cl electrodes that were placed on the scalp according to the international 10/20 system. The data were digitized at 5 kHz, referenced online to FCz using a non-magnetic MRI-compatible EEG system (BrainAmp MR plus, Brain products, Munich, Germany). Impedances were kept below 10 k before recording. Electrocardiogram (ECG) was simultaneously acquired from each participant. The EEG amplifier used a rechargeable power pack that was placed outside the scanner bore. To ensure the temporal stability of the EEG acquisition in relation to the switching of the gradients during the MR acquisition, a SyncBox (SyncBox MainUnit, Brain Products GmbH, Gilching, Germany) was used to synchronize the amplifier system with the MRI scanner’s system. Fiber optic cables transmitted the amplified and digitized EEG signal to the recording computer, which was outside the scanner room. An Adapter (BrainAmp USB-Adapter, Brain products, Gilching, Germany) was used to convert optical into electrical signal.

#### fMRI

All images were acquired using a 3T Siemens Trio scanner. A T2-weighted gradient echo planar imaging (EPI) sequence (TR/TE=1500/29 ms, FOV=192 × 192 mm2, flip angle=90 deg, acquisition matrix=64 × 64, thickness/gap=5/0.5 mm, in-plane resolution=3.0 × 3.0 mm2, axial slices=25) was used for functional image acquisition. The first three volumes of each sequence were discarded to account for magnetization saturation effects. All subjects were scanned in six blocks, where each block lasted 435 s and contained 290 volumes. After the first three blocks, a T1 was acquired for 5 min, where participants were told to relax and hold still. The 3D spoiled gradient recalled (SPGR) sequence used the following parameters: TR/TE=8.5/3.4 ms, FOV=240 × 240 mm2, flip angle=12 deg, acquisition matrix=512 × 512, thickness=1 mm with no gap. The high-resolution T1-weighted structural volume provided an anatomical reference for the functional scan. We minimized head movements by using a cushioned head fixation device.

### Behavioural Data Analysis

Recall accuracies at the final memory test were estimated for Think, No-Think and Baseline conditions, and for each test type (SP and IP) separately. The analyses were conditionalized: only word pairs learned in the study phase (determined by the memory test prior the TNT phase) were considered. Recall accuracies were computed as a ratio between the number of word pairs correctly recalled at the final test and the total number of word pairs that were learned at study. These measures were compared using a two-way analysis of variance (ANOVA) with the memory condition (No-Think and Baseline) and test type (SP and IP) as factors, to determine whether there was below-baseline forgetting. Paired-samples T-tests were applied to assess the effect of memory suppression and memory retrieval on final recall performance within each test type. We conducted all behavioural statistical analyses in SPSS Statistics 19.0.

### EEG data preprocessing

Main fMRI gradient and ballistocardiogram (BCG) artifacts were first removed from the EEG data acquired during the TNT phase, following standard template subtraction procedures. Subsequently, data were down-sampled to 250 Hz and digitally filtered within 0.1-45 Hz using a Chebyshev II-type filter. Temporal independent component analysis (ICA; Bell and Sejnowski, 1995) was subsequently applied to attenuate ocular artifacts (e.g., blinks, saccades), fronto-temporal muscular activity of small intensity and residual BCG and imaging artifacts. Artifactual components were visually selected on the base of their characteristic time courses, topographic amplitude distributions, signal features (i.e., kurtosis, energy) and spectral characteristics (Mayeli et al., 2016). Continuous EEG data was divided into 5000-ms segments relative to the onset of all words presented during the TNT phase (Think, No-Think and Perceptual Baseline). Each segment included 500 ms of pre-stimulus baseline and 4500 ms of post-stimulus period. Segments that were contaminated by jumps, movement or strong muscular activity were removed. Finally, EEG signals were re-referenced to the average for further analyses. All following analyses were conditionalized as the behavioural measures by including only trials belonging to word pairs that were learned at study.

### ERP analyses

ERPs were computed within the 1250-ms epoch comprising 250 ms of pre-stimulus baseline and 1000 ms of post-stimulus period. The suppression-N2 component was identified by visual inspection of the grand-mean ERPs of frontocentral sensors, around the latencies reported in previous studies (Mecklinger et al., 2009). Statistical comparisons between No-Think and Think N2 waveforms were limited to the 300-450 ms time window. A first analysis contrasted the mean amplitudes of all sensors in two overlapping 100-ms windows using non-parametric paired T-tests. Cluster-based permutation tests with 5000 Monte Carlo randomizations were applied to correct for multiple comparisons across time and sensors. Each iteration assigned random conditions labels to each trial and extracted the cluster of sensors (p*<*0.05, one-tailed) with maximal (negative) summed statistic. To determine the precise time window of the N2 effect, a second analysis contrasted amplitudes from all time points of a pooled frontocentral channel (Fz, FC1 and FC2). To correct for multiple comparisons across time points, another permutation test with 5000 Monte Carlo randomizations was applied, based on the maximal (negative) statistic. To investigate the relationship between the N2 effect and forgetting, robust Pearson correlations were computed between the differential waveform of the pooled channel (mean Think minus No-Think amplitude within the significant window) and the mean SIF in both memory tests.

### Time-frequency analyses

Time-frequency representations (TFRs) were computed on data-padded wider epochs (−3000-5500 ms) to prevent edge filter effects. Epochs were convolved with 6-cycles and 3-cycles wavelets and then cropped to obtain 2-30 Hz spectral power between -500 ms pre-stimulus to 3000 ms post-stimulus, in 50 ms by 1 Hz time-frequency bins. To further reduce the contribution of noise, participants TFRs were normalized using a single-trial baseline correction method (Grandchamp and Delorme, 2011). First, power values of each time-frequency bin and channel were z-transformed using the mean power and standard deviation across all trials. After trial average, TFRs were converted into z-power change relative to baseline (−500 ms to 0 ms pre-cue window) by subtracting the mean baseline z-power value from all time points of each frequency bin and channel. Within-condition relative power increases or decreases were determined by contrasting each time point against the mean baseline value at each frequency bin using non-parametric paired T-tests. P-values were computed through 5000 Monte Carlo randomizations. Then, false discovery rate (FDR) procedure (Benjamini and Hochberg, 1995) was applied (p*<*0.05) to correct for multiple comparisons across time-frequency bins and scalp locations. TFRs were contrasted between Think and No-Think conditions using cluster-based permutation tests with 5000 Monte Carlo randomizations to correct for multiple comparisons across time-frequency bins and scalp locations. Each iteration assigned random conditions labels to each trial and extracted the cluster of sensors and time-frequency bins (p*<*0.05) with maximal summed statistic. A similar statistical procedure was followed to compare TFRs between large and small N2 trials.

### Source localization

We created realistic 3-shell boundary element models (BEM) based on individual T1 MRIs. Each BEM consisted of 3 closed, nested compartments with conductivities .33 S/m, .0042 S/m and .33 S/m corresponding to skin, skull, and brain, respectively. To obtain the models, skull and brain binary images were obtained using Fieldtrip segmentation routine. Scalp voxels were first identified with the SPM8’s “New Segment” algorithm (Ashburner and Friston, 2005) combined with an extended tissue probability map (eTPM) that includes eyeballs, the whole head and the neck (for more details, see procedure and code provided by Huang et al., 2013). Resulting scalp probability maps (including eyeballs) were smoothed and binarized. All binary images were manually corrected using MRI visualization software to fit better the anatomy and make them suitable for BEM computation (i.e., remove overlaps and irregularities). It was particularly critical to correct the scalp masks because EEG sensors were automatically classified as scalp tissue and would have produced bumpy models otherwise. Binary masks were used to create boundary meshes in Fieldtrip using iso2mech method with 10000 vertices, which were smoothed afterwards. Real EEG sensor coordinates were determined by hand from rendered models of raw scalp masks using MRI visualization software (ITK-SNAP). This was possible because EEG sensors were visible in the T1s and appeared as small bumps on the rendered models. TP9 and TP10 sensors (behind the ears) were often difficult to identify and were excluded from source localization analyses to reduce localization errors. A grid of source locations was defined in individual’s brain space but corresponding to 1-cm grid MNI coordinates that were consistent across participants. To obtain individual coordinates, MRIs were normalized to the standard T1 template, using non-linear transformations in SPM12. The inverse of this transformation was then applied to the template source grid obtained in Fieldtrip. The leadfields were computed with OpenMEEG v2.4 (Gramfort et al., 2010) called from Fieldtrip, using international units (i.e., EEG amplitudes in Volts and electrode and source-grid coordinates in meters).

To localize ERP and TFRs effects, we employed linearly constrained minimum variance (LCMV) beamformer (Van Veen et al., 1997). To enhance the detection of N2 sources from less superficial brain areas such as those in dACC, we first subtracted the trial-averaged data (0.5-30 Hz) of Think from No-Think condition. This method is called subtraction of epoch-averaged sensor data (SAD), and is one of the subtraction techniques recommended to reduce interference from dominant sources common to two experimental conditions (Mills et al., 2012). The spatial filters were first computed within the time window showing significant amplitude differences at sensor-level (300-570 ms), and then applied to the N2 and baseline time windows to obtain a power distribution map per participant. Participants’ power distribution maps entered a group-level one-sample T-test against zero. To correct for multiple comparisons, we applied the maximal statistic method using 5000 Monte Carlo randomizations implemented in Fieldtrip (Oostenveld et al., 2011). We used a similar procedure to localize the sources within the time window of maximal amplitude differences in dACC ROI (548-708 ms). To determine whether source activity differences were task-related, power distribution maps were contrasted with baseline power maps using paired-sample T-tests and cluster-based statistics with 5000 randomizations. To determine the time window of maximal amplitude differences within dACC ROI, dipole momentum time-courses were extracted for each cartesian direction and averaged across trials for each condition. Think and No-Think amplitudes were compared with paired-sample T-tests and cluster-based statistics as described for sensor-level ERP analyses, to correct for multiple comparisons across time points, source locations and orientations. To localize TFR effects, the spatial filters were computed over the whole epoch (−500-3000 ms) without averaging. For each pre-defined source location, dipole momentum time-courses were extracted for each cartesian direction and collapsed into a single trace after determining the principal component. TFRs were computed and z-transformed as described for sensor-level. For statistical analyses, power values within the time-frequency window of interest were averaged and translated to brain maps. Contrasts were performed with paired-sample T-tests and cluster-based statistics as for ERP analyses. For ROI analysis, TFRs from all voxels within an ROI were identified and averaged across trials for each condition. Think *Crespo-García, Wang, Jiang, Anderson, and Lei* Anterior Cingulate Cortex Signals *Preprint v1*.*1, Aug*.*06, 2021* and No-Think amplitudes were compared with paired-sample T-tests and cluster-based statistics, to correct for multiple comparisons across time-frequency bins and source locations. The regularization parameter was set at 0.001 % of the largest eigenvalue of the covariance matrices.

### ROIs definition

To test our a-priori hypotheses, some analyses were restricted to the dACC, rDLPFC and bilateral hippocampus. dACC, and rDLPFC masks only included subregions that are commonly activated during action cancellation and memory inhibition, revealed by meta-analyses using fMRI data from stop-signal and TNT tasks (Apšvalka et al., 2020; Guo et al., 2018). Each mask comprised the cluster of voxels revealed by conjunction analyses combining the contrasts No-Think*>*Think and Stop*>*Go (e.g., Figure 3.2 of Guo et al., 2018). Left and right hippocampal masks were constructed from a probabilistic map based on cytoarchitectonic delimitations derived from 10 post-mortem human brains warped to the MNI template brain (Amunts et al., 2005). These maps contain the relative frequency with which a cerebral structure was present on each voxel of the anatomical MNI space. A binary mask was built from the region having a probability of 0.1 or higher to be labelled as hippocampal (CA, DG, Subc) or entorhinal cortex (EC).

### Granger causality analyses

To investigate the effective connectivity and directionality of the information flow between our a-priori selected ROIs, we performed Granger causality (GC) analyses on EEG source activity. We computed spectral GC (Geweke, 1982) estimates within the 1-s time window right after the N2 (450-1450 ms) using the Fourier-based nonparametric approach (Dhamala et al., 2008) as implemented in Fieldtrip (Oostenveld et al., 2011). We chose the nonparametric instead the autoregressive approach because it does not require to determine the optimal model order for each participant. The window width was defined to be relatively short as to fulfil the stationarity assumption and have some temporal resolution, long enough to include sufficient data, and has been used in a previous application of the nonparametric approach in a similar context (Popov et al., 2018). To further approximate stationarity, we subtracted the event-related potential from the data before computing the Fourier transform for the whole spectrum (frequency smoothing of 2 Hz). Thereafter, the Fourier coefficients were used to compute the cross-spectral density matrix. This matrix was then factorized into the noise covariance matrix and the spectral transfer matrix which are necessary for calculating GC (see Dhamala et al., 2008 for more details). To determine the statistical significance of the directionality of information flow between each pair of sources, we compared the magnitude of Granger coefficients (2-30 Hz) for both possible directions (from source 1 to source 2 and vice versa) and trial types (e.g., Think and No-Think) using cluster-based permutation ANOVAs and T-tests. In addition, to confirm that the observed differences in GC values were caused by true directed relationships and not by differences in signal-to-noise ratio (SNR), we computed GC on the time-reversed source activities. After doing this control analysis, true causal interactions should show an inversion in the directionality of the information flow, whereas spurious interactions would appear as unchanged.

### fMRI data preprocessing

FMRI data were preprocessed using SPM12 software (Henson et al., 2019). Standard preprocessing steps were applied, including: (i) spatial realignment to correct for head movements, (ii) slice timing, (iii) coregistration of the structural to the functional images, (iv) segmentation of the coregistered structural image (which performs spatial normalization to MNI space and generates a deformation field file), (v) normalization of the functional images applying the deformation field, and (vi) spatial smoothing with a three-dimensional 6 mm full-width at half maximum Gaussian kernel. As an additional control for head-movement artifacts, we excluded two data blocks in a participant because its mean head-movement regressors exceeded 4 mm in one of the orthogonal directions.

### fMRI data analysis

FMRI data acquired during the TNT phase was analysed through general linear models (GLMs) in SPM12. Each data block for a given participant was modelled separately at the first level using a fixed-effects model. Then, each group analysis used a random-effects model. To identify the regions that were engaged or downregulated during retrieval suppression, we constructed a GLM containing three regressors of interest and one regressor of no interest. Regressors of interest were built with the onset times of the words presented in the three experimental conditions (Think, No-Think and Perceptual Baseline). fMRI analyses were conditionalized, so Think and No-Think onset times corresponding to word pairs that participants failed to learn at study (Misses) were excluded from the main regressors and grouped in the regressor of no interest. Six further regressors contained the head-movement parameters obtained during spatial realignment and were included as covariates. First-level analyses calculated the main effects between the three conditions. For group-level analyses, contrasts between No-Think and Think, each relative to Perceptual baseline, were compared by means of paired-sample T-tests. For whole brain analyses, activations were considered significant if formed by clusters of at least 20-voxels with an uncorrected p-value smaller than 0.001. For dACC ROI analysis, a paired-sample t-test was applied to averaged contrast-values (p*<*0.05).

### ERP-informed fMRI analyses

To determine whether and how (e.g., positive or negative modulation) ROI’s BOLD responses covaried with the EEG measures across trials, we chose a parametric design approach (Debener et al., 2006). For each EEG measure, we built a separate GLM. Each GLM extended the previously described GLM by including two additional regressors of interest and one of no interest. The regressors of interest were parametric modulator vectors for No-Think and Think conditions containing single-trial values of the corresponding EEG measure. The regressor of no interest gathered the onset times of all trials that were classified as artifacts during EEG pre-processing or that showed EEG measures larger or smaller than 3 standard deviations (i.e., these onset times were removed from the original main regressors). One of the parametric modulators was built from single-trial N2 amplitude values from the pooled frontocentral channel in the selected time window. To extract single-trial N2 amplitudes, an individual search window was first limited to participants P2 and P3 waveforms of the pooled channel ERP. Then, N2 latencies were defined as the minimum peak found within the individual search window for each trial. To help the peak detection algorithm, data were low pass filtered at 8 Hz. However, single-trial N2 amplitudes were obtained from the 0.5-30 Hz data, by averaging 100-ms windows centred on the N2 latencies. A second parametric modulator was built from mean theta (4-8 Hz) power values extracted from dACC ROI in the N2 time window. Other parametric modulators (i.e., hippocampal ROIs theta power) were built from mean power values across all voxels within a cluster. We computed other GLMs for control purposes. In one case, we extracted the mean dipole momentum from dACC ROI within the N2 effect time window. To control for the sign of the source time-courses and the correlation with BOLD, we run three separated GLMs, one for each cartesian direction, and the resulting contrasts were averaged. The signs of the source time-courses followed the same convention of the frontocentral channel in that window (i.e., more negative for larger N2 amplitudes, compared to small N2 amplitudes). For group-level analyses, contrasts of parameter estimates were averaged across all voxels within each ROI for each condition and subject. Within-condition and between-condition contrasts were assessed using one-sample and paired-sample and independent-sample T-tests, respectively. Whole-brain statistical maps were corrected for multiple comparisons using a cluster-based permutation approach. For each permutation, we used the same original GLM, but the order of the values in the parametric modulator (i.e., amplitude or latency) was randomized. As a result, the new parametric modulators contained the same values than the original regressor, but each value was assigned to an onset time of a different trial. This procedure was repeated 100 times in all participants (first-level fixed effects), which yielded 100 second-level fixed effects analyses. We recorded the sizes of all clusters obtained from the resulting statistical maps after applying an initial threshold at |Z|score*>*2.57 (uncorrected p*<*0.005). Then, we generated a distribution with the cluster sizes, which enabled us to determine the largest cluster size leading to a significance level of p*<*0.05. This cluster size was used as a threshold to correct the original statistical maps (Fouragnan et al., 2017).

## Acknowledgments

We thank Xu Xiaoxiao, Dong Debo and Yang Tianliang for their assistance in conducting the EEG-fMRI experiment, Dace Apšvalka for providing the meta-analysis ROIs and advice on fMRI data quality and univariate analyses, Richard Henson and Johan Carlin for key advice on EEG-informed fMRI analyses, Davide Nardo for comments on a draft of this manuscript and advice on fMRI statistics, Haakon Engen for training on fMRI data preprocessing, and to Daniel Wong and Sarang S. Dalal for useful scripts supporting the construction of model meshes. This research was supported by grants from the National Natural Science Foundation of China (31971028 to X.L.), Major Project of Medicine Science and Technology of PLA (AWS17J012 to X.L.), and Royal Society’s Newton International Fellowship (NF171519 to M.C.G.).

## Author Contributions

M.C.A. and X.L. conceived the experiment. Y.W. performed the experiment. M.C.G., Y.W., X.L. and M.J. analysed the data. M.C.G. and M.C.A wrote the manuscript. M.C.A., X.L., Y.W. and M.J. discussed the results and contributed to the manuscript.

