## Supplementary data for "Anterior Cingulate Cortex Signals the Need to Control Intrusive Thoughts During Motivated Forgetting"

### Supplemental Results

It is well known that motion can introduce artifacts in EEG signals recorded inside the magnetic field of an MR scanner. Moreover, if motion correlates with experimental manipulations, it could cause spurious task-related EEG power effects as well as spurious EEG-BOLD correlations (Fellner et al., 2016). We did not find evidence that our results were affected by this problem. First, sensor-level oscillatory power effects resulting from contrasting No-Think and Think conditions agreed well with previous studies using out-of-scanner EEG data. Second, we did not find significant between-condition differences in motion measures (Figure S6). Third, correlations between oscillatory power and motion measures were minimal and could not explain observed power differences between conditions (Figure S7). Fourth, power-BOLD correlations were performed using source activity obtained with beamforming, not sensor activity. Beamforming is a spatial filter and therefore it works as a denoising method because it can separate motion activity generated around the sensors from deeper neural activity. Fifth, correlations between BOLD signal and motion measures during memory suppression were negative in bilateral putamen, cerebellum, left inferior parietal lobe and medial PFC, including regions of dACC. However, these effects did not overlap the areas showing significant correlations between BOLD signal and sensor EEG measures (Figure S8A). Furthermore, positive correlations between BOLD signal in dACC ROI and frontocentral N2 amplitudes were enhanced when including motion measures as additional regressors (new GLM: t-max = 2.91, $p = 0.004$; original: t-max = 2.46, $p = 0.011$). In addition, N2-BOLD signal correlations derived from GLM models including motion measures as additional regressors were significant in the same set of brain areas derived from the original GLM models (Figure S8B). Correlations between motion and BOLD signals were only more negative in left cerebellum during memory suppression, compared to retrieval.

### Supplemental Material and Methods

We performed several control analyses to inspect whether power-BOLD coupling effects were driven by motion artifacts that could have remained in the EEG data. We contrasted No Think and Think TFRs and compared the effects to those reported in previous literature. Statistical analyses were performed using a cluster-based permutation test as described in the STAR Methods.

#### *Motion measures*

Movement measures were computed from the 6 head-movement parameters obtained during spatial realignment (see methodological details in (Fellner et al., 2016)). First, each head-movement parameter was normalized using its mean and standard deviation computed over all scans. Then, a motion measure was calculated by subtracting the normalized head-movement parameter at scan *s* from that of the subsequent scan (*s* + 1) and squaring the resulting values. The 6 motion measures were averaged and normalized to obtain a final z-transformed motion measure that was used in the following analyses. To investigate possible differences in motion between experimental conditions, stimulus onsets during the TNT phase were temporally matched to fMRI scans. Each stimulus onset inherited the motion value of the closest scan. Mean z-motion measures were computed for each condition and participant and compared between conditions using paired t-tests at group-level.

#### *EEG power and motion*

The relationship between oscillatory power and motion was investigated by a two-level correlation analyses(Fellner et al., 2016). First, Spearman's correlations between TFRs of all artifact-free TNT epochs and motion values were computed within participant, for each electrode and time-frequency bin. The resulting correlation coefficients were Fisher z-transformed and entered the group-level analysis. Z-correlation coefficients were compared against zero using Wilcoxon sign-ranked tests. To correct for multiple comparisons across time, frequency and electrodes, a cluster-based permutation approach with 5000 Monte Carlo randomizations was applied. Each iteration assigned random labels of task and zero conditions to each trial and extracted the cluster of sensors ($p < 0.05$) with maximal summed statistic.

***BOLD and motion***

To inspect correlations between BOLD signals and motion, we constructed new GLMs like those used for EEG-informed fMRI analyses with EEG measures. However, in this case, z-motion measures were introduced as parametric modulators and convolved with the hemodynamic response function (HRF). To identify brain regions showing pure correlations with z-motion measures, a single parametric modulator was introduced in a version of the analyses. To verify that correlations between sensor-level EEG measures (frontocentral N2 amplitudes and FMT power) were not caused by motion, we performed new EEG-informed fMRI analyses with EEG measures as regressors of interest but adding a second regressor that included z-motion measures orthogonalized relative to the task regressors. The z-motion measures were also convolved with the hemodynamic response function (HRF). Statistical analyses were performed like for the main EEG-informed fMRI analyses with EEG measures.

### Supplemental Figures

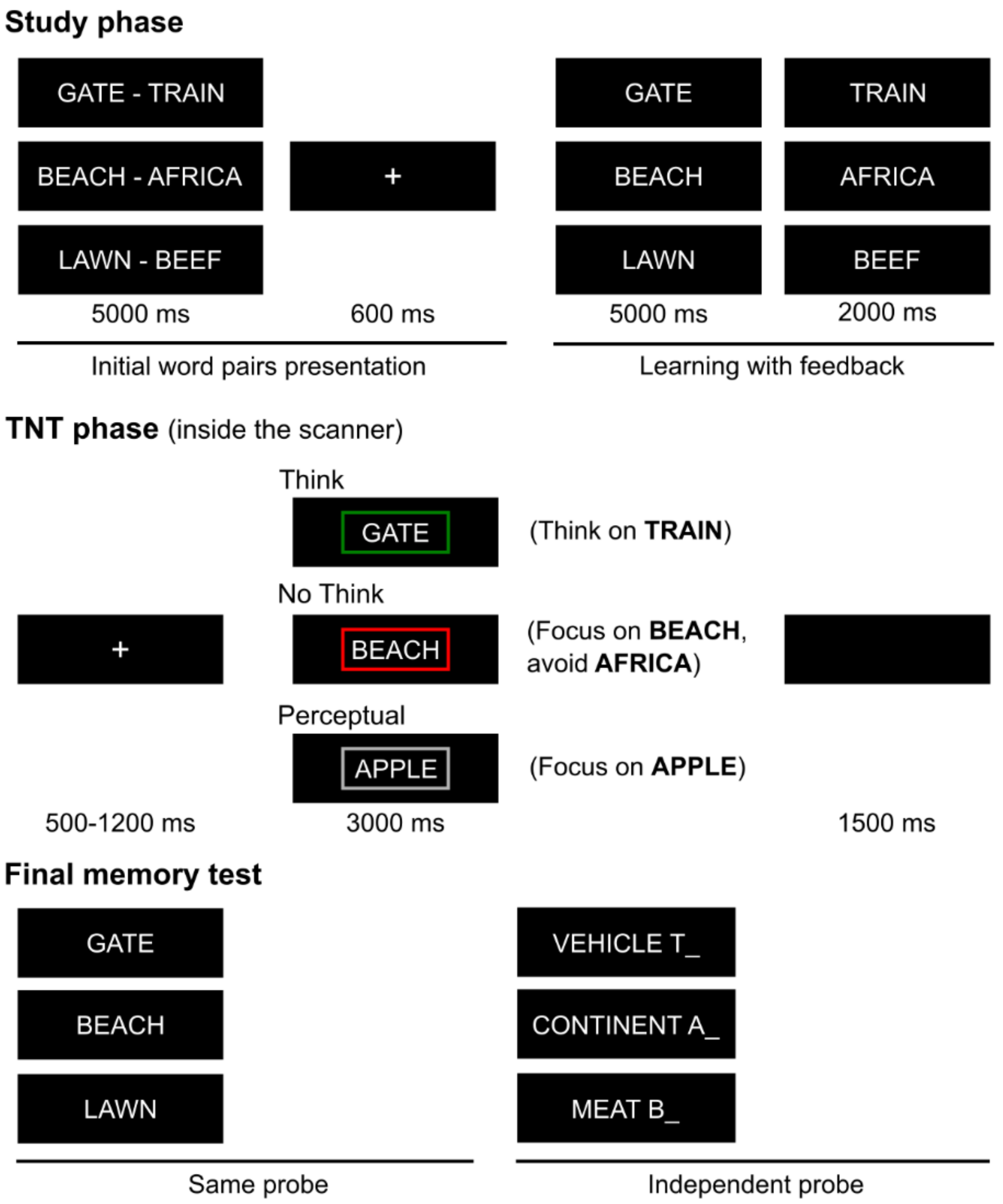


**Figure S1. Experimental paradigm**

In the study phase, participants encoded cue-associate word pairs. During the scanned TNT phase, participants recalled some of the associates (Think, cues presented inside a green frame) and suppressed others (No-Think, cues presented inside a red frame). Participants payed full attention to unpaired words (without associate) presented inside a grey frame (Perceptual). In the final memory phase, participants were asked to recall the associates given their cues (same probe) or their category name (independent probe). Memory was also evaluated for some word pairs that were initially learned but did not enter the TNT phase (Baseline).

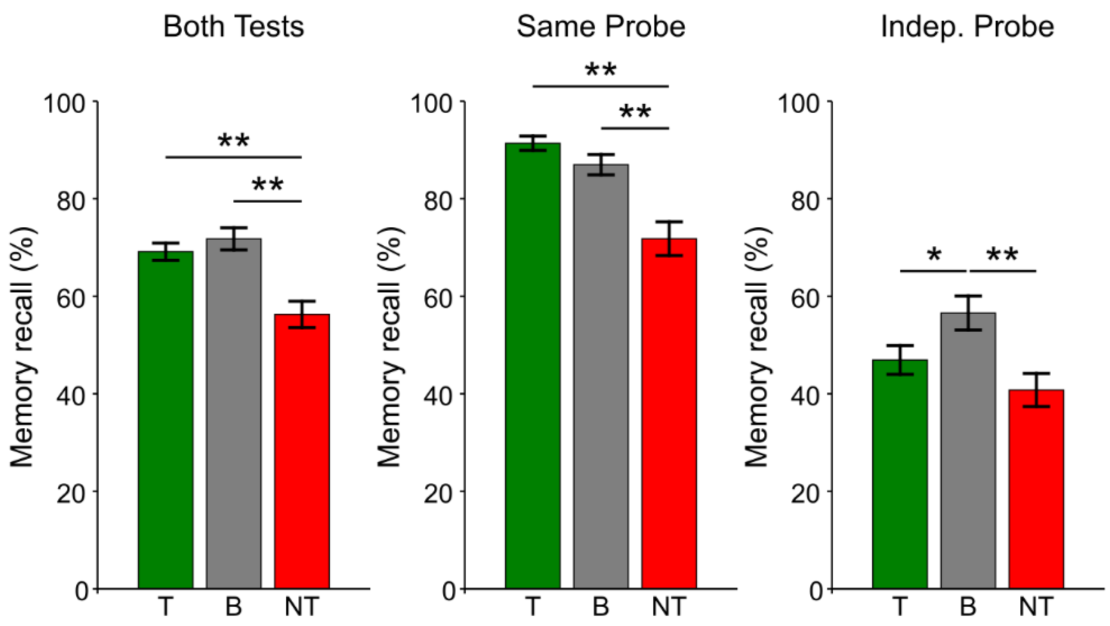


**Figure S2. Behavioural results**

Percentage of items correctly recalled in the final memory tests using same probes (middle panel) and independent probes (right panel). Mean values across both tests are shown in left panel. Analyses were conditionalized on participants' recall before entering the TNT phase. Error bars show ± 1 standard error of the mean. Stars denote significant effects: ** p<0.001, * p<0.05. T = Think, B = Baseline, NT = No-Think.

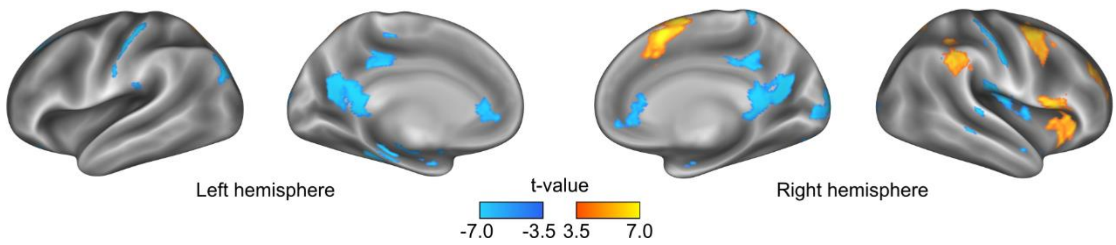


**Figure S3. fMRI contrast No-Think vs Think**

Whole-brain paired-sample contrast between No-Think and Think conditions. Hot colors indicate brain areas more active during retrieval suppression than during retrieval (No Think > Think), whereas cold colors indicate the opposite (No Think < Think). T-maps show values with p < 0.001 (uncorrected) and clusters with more than 20 voxels.

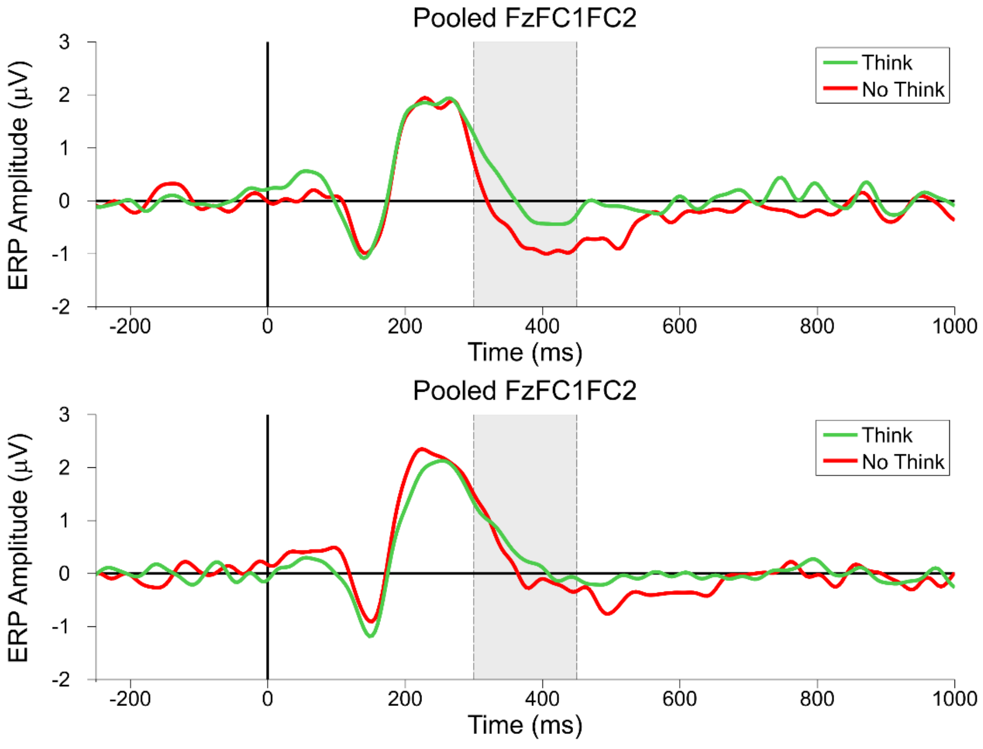


**Figure S4. ERP analyses in high and low forgetters**

High forgetters showed N2 effect (top panel) whereas low forgetters did not (bottom panel). ERP waveforms for the Think (green) and No Think (red) conditions for the frontocentral channel. Gray shadow indicates the time window considered for N2 wave analyses (300-450 ms).

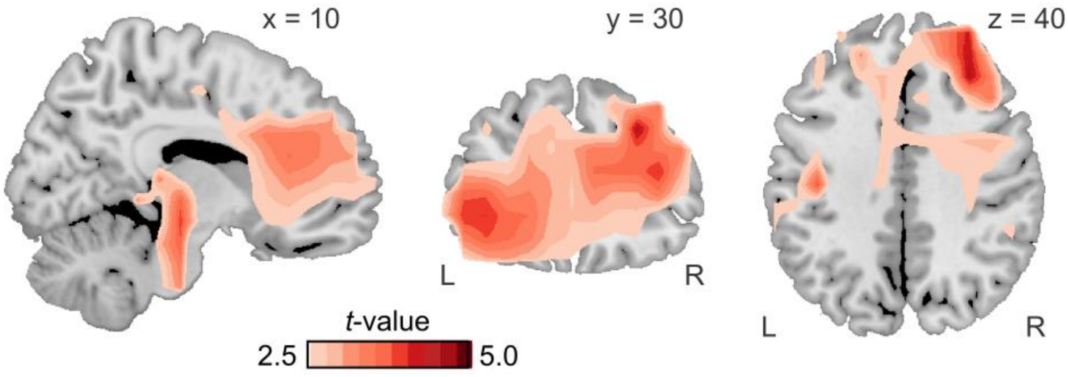


**Figure S5. Sources of No-Think vs Think effect in the delayed control window**

Source maps showing above-baseline No-Think vs Think effect in evoked activity during the delayed control window (0.5-8 Hz; 428-728 ms) over frontal regions, including dACC and rDLPFC ($p < 0.05$, cluster-based correction).

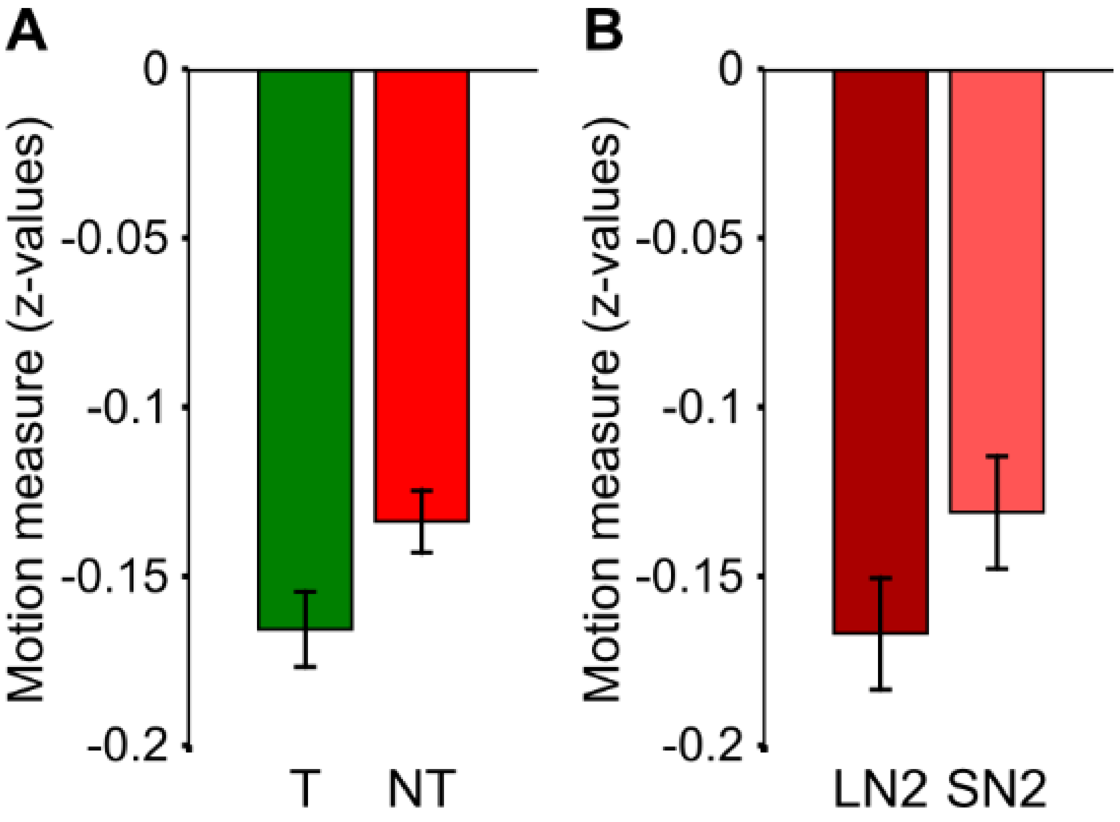


**Figure S6. Bar plots of mean motion in the TNT phase**

(A) There were not significant differences in motion between Think and No-Think trials ($t(23) = -1.76$, $p = 0.09$). (B) There were not significant differences in motion between No-Think trials with large N2 and small N2 No-Think trials ($t(23) = -1.16$, $p = 0.26$). Error bars show ± 1 standard error of the mean.

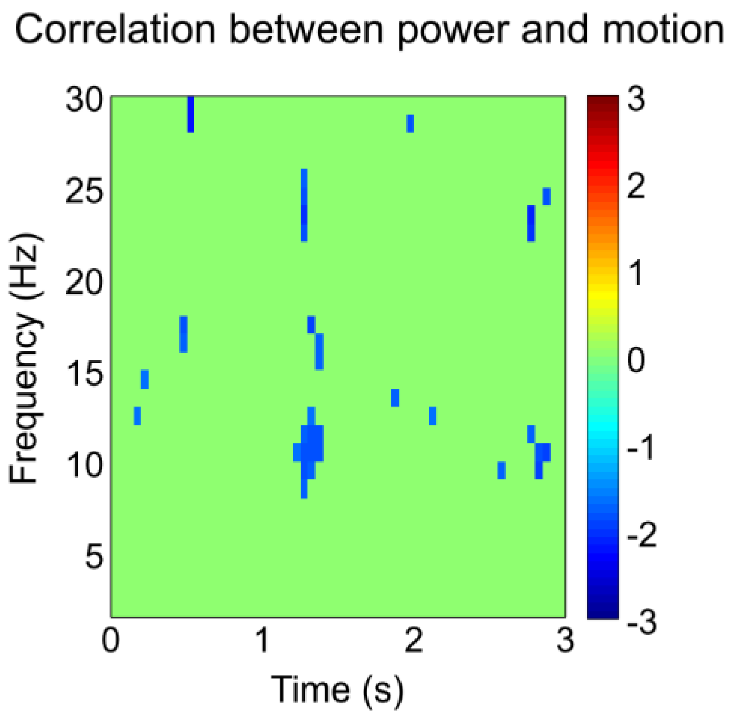


**Figure S7. TFR power correlation with motion**

Time-frequency representation (TFR) map showing negative correlations between task oscillatory power and motion measures. To highlight significant time-frequency bins surviving cluster-based permutation test correction ($p < 0.05$), non-significant bins are masked in green.

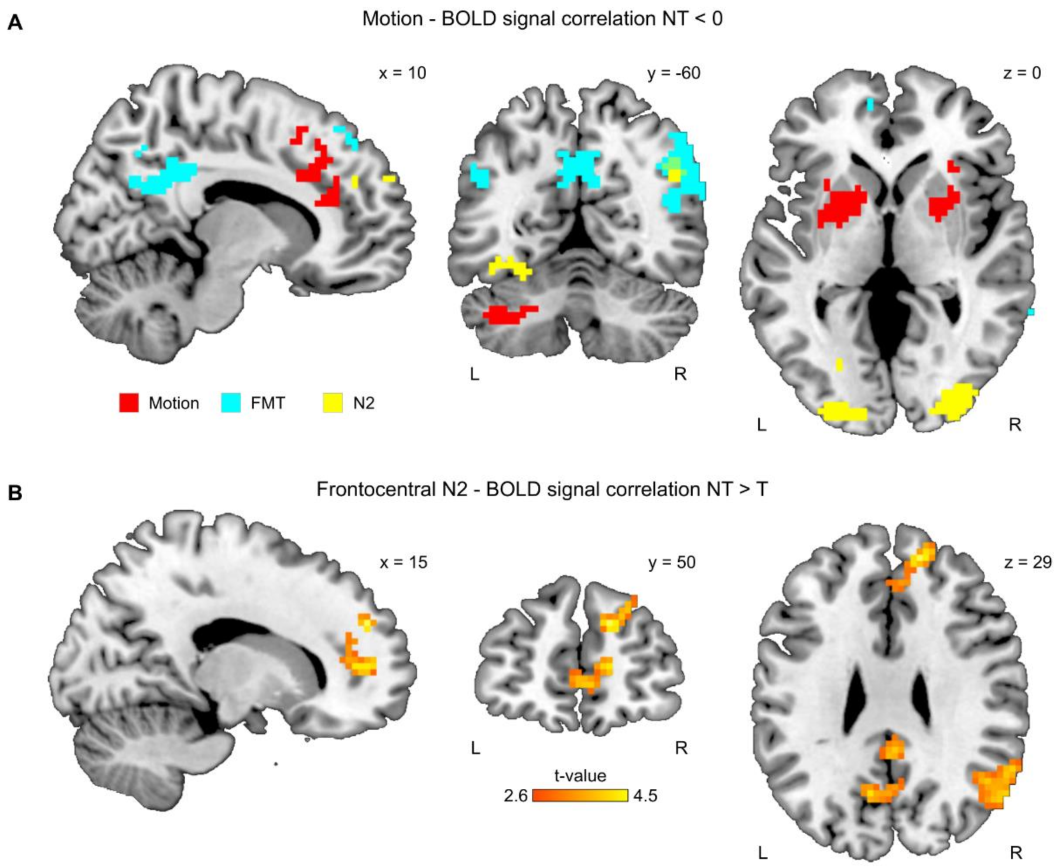


**Figure S8. Motion–BOLD signal correlations**

(A) Brain areas showing negative motion-BOLD signal correlations during memory suppression (red; cluster extent >= 35) do not overlap with brain areas showing positive correlations with N2 amplitudes (yellow; cluster extent >= 35) or negative correlations with FMT (cyan; cluster extent >= 40). All uncorrected $p < 0.005$. (B) The set of brain areas showing stronger N2-BOLD signal correlations during suppression relative to retrieval remain the same after correcting for motion (motion measures were included as an additional regressor in the GLM).

### Supplemental Tables

**Table S1. Overview of fMRI Results**

| Contrast and Brain Regions | ~BA | MNI Coordinates x | y | z | Z Score | T Score | Cluster extent | p-cluster value |
|---|---|---|---|---|---|---|---|---|
| **No-Think > Think** | | | | | | | | |
| **Right medial SFG/SMA** | 6 | 9 | 11 | 59 | 5.66 | 8.49 | 293 | < 0.001 |
| **Right supramarginal gyrus** | 40 | 60 | -46 | 38 | 4.78 | 6.35 | 174 | < 0.001 |
| **Right MFG/precentral gyrus** | 6 | 30 | -1 | 47 | 4.72 | 6.23 | 223 | < 0.001 |
| **Right IFG/insula** | 45 | 54 | 17 | 5 | 4.52 | 5.83 | 299 | < 0.001 |
| **Right SPL** | 7 | 21 | -61 | 59 | 4.41 | 5.61 | 95 | 0.010 |
| **Right MFG** | 9/10 | 36 | 38 | 32 | 4.02 | 4.90 | 76 | 0.025 |
| **Left SPL/precuneus** | 7 | -21 | -67 | 62 | 4.00 | 4.89 | 64 | 0.047 |
| **Think > No-Think** | | | | | | | | |
| **Left PCC/precuneus** | 23/29/30 | -9 | -49 | 14 | 5.43 | 7.86 | 482 | < 0.001 |
| **Left hippocampus** | 54 | -33 | -34 | -10 | 4.84 | 6.48 | 96 | 0.009 |
| **Left MFG/SFG** | 6/8 | -30 | 23 | 59 | 4.61 | 6.00 | 119 | 0.003 |
| **Right postcentral gyrus/insula** | 13 | 45 | -22 | 23 | 4.59 | 5.97 | 156 | 0.001 |
| **Right postcentral gyrus** | 6/4 | 54 | -10 | 56 | 4.54 | 5.87 | 116 | 0.004 |
| **Right postcentral gyrus** | 2 | 15 | -46 | 77 | 4.52 | 5.82 | 83 | 0.018 |
| **Left putamen** | | -12 | 2 | -10 | 4.51 | 5.81 | 63 | 0.050 |
| **Right cingulate cortex/SMA** | | 6 | -40 | 38 | 4.45 | 5.69 | 183 | < 0.001 |
| **Right occipital cortex/FG** | 18/17 | 18 | -94 | 11 | 4.39 | 5.57 | 237 | < 0.001 |
| **Left postcentral gyrus** | 4/6 | -60 | -7 | 38 | 4.31 | 5.43 | 96 | 0.009 |
| **Left ACC** | 32 | 0 | 41 | 5 | 4.21 | 5.23 | 253 | < 0.001 |
| **Left angular gyrus** | 39/19/40 | -39 | -79 | 32 | 3.88 | 4.67 | 104 | 0.006 |

**BA, Brodmann's area; ACC, anterior cingulate cortex; IFG, inferior frontal gyrus; FG, fusiform gyrus; MFG, middle frontal gyrus; PCC, posterior cingulate cortex; SFG, superior frontal gyrus; SMA, supplementary motor area; SPL, superior parietal lobe. Cluster extents are indicated in number of voxels. p-cluster values are corrected for multiple comparisons using familywise error (FWE), p(FWE) < 0.05. Initial p-voxel threshold at p < 0.001 uncorrected.**

| **Table S2. Coupling N2-theta power dACC source and BOLD signal** | | | | | | | | |
|---|---|---|---|---|---|---|---|---|
| | | MNI Coordinates | | | | | | |
| **Contrast and Brain Regions** | ~BA | x | y | z | Z Score | T Score | Cluster extent | p-uncorrected |
| **No-Think < 0** | | | | | | | | |
| **Right precuneus/PCC** | 7/31 | 6 | -58 | 44 | 3.84 | 4.60 | 532 | < 0.001 |
| **Left medial SFG** | 9/10 | -3 | 44 | 26 | 3.79 | 4.52 | 253 | < 0.001 |
| **Right lateral/medial SFG** | 10 | 30 | 59 | 20 | 3.72 | 4.42 | 180 | < 0.001 |
| **Left medial orbital PFC/ACC** | 24 | 0 | 50 | -16 | 3.57 | 4.18 | 65 | < 0.001 |
| **Right STG** | 39 | 48 | -55 | 20 | 3.52 | 4.10 | 50 | < 0.001 |
| **Left insula/inferior/orbital PFC** | 13/47 | -24 | 17 | -13 | 3.49 | 4.06 | 88 | < 0.001 |
| **Right IPL** | 39/40 | 42 | -70 | 41 | 3.33 | 3.82 | 75 | < 0.001 |
| **Right MTG** | 21 | 57 | -37 | -13 | 3.24 | 3.69 | 120 | < 0.001 |
| **Left putamen** | | -33 | 2 | -4 | 3.21 | 3.66 | 51 | < 0.001 |
| **Right putamen** | | 27 | -13 | 2 | 3.00 | 3.36 | 48 | < 0.001 |
| **No-Think < Think** | | | | | | | | |
| **Right PCC** | 31 | 3 | -28 | 41 | 4.30 | 5.40 | 296 | < 0.001 |
| **Left orbital/superior medial PFC** | 11 | 0 | 53 | -13 | 4.09 | 5.03 | 460 | < 0.001 |
| **Left parahippocampus/hippocampus** | 36 | -9 | -40 | -25 | 4.08 | 5.02 | 50 | < 0.001 |
| **Right MFG/SFG** | 6 | 39 | -4 | 59 | 3.98 | 4.85 | 47 | < 0.001 |
| **Left SFG/MFG** | 8 | -15 | 38 | 50 | 3.96 | 4.81 | 101 | < 0.001 |
| **Right SFG** | 10 | 27 | 59 | 17 | 3.96 | 4.81 | 45 | < 0.001 |
| **Left cerebellum** | | -42 | -64 | -46 | 3.66 | 4.32 | 111 | < 0.001 |
| **Right middle temporal gyrus** | 21/22 | 66 | -46 | -1 | 3.64 | 4.29 | 63 | < 0.001 |
| **Left PCC/ Precuneus** | 23 | -6 | -55 | 8 | 3.56 | 4.16 | 52 | < 0.001 |
| **Left AG/MTG/occipital cortex** | 39 | -45 | -67 | 44 | 3.54 | 4.13 | 99 | < 0.001 |
| **Left MTG/temporal pole** | 38 | -54 | 11 | -28 | 3.48 | 4.04 | 41 | < 0.001 |
| **Right MTG/temporal pole** | 21 | 57 | -4 | -22 | 3.35 | 3.86 | 46 | < 0.001 |

**BA, Brodmann's area; ACC, anterior cingulate cortex; AG, angular gyrus; IPL, inferior parietal lobe; MFG, middle frontal gyrus; MTG, middle temporal gyrus; PCC, posterior cingulate cortex; PFC, prefrontal cortex; SFG, superior frontal gyrus; STG, superior temporal gyrus. Cluster extents are indicated in number of voxels. Initial p-voxel threshold at p < 0.005 uncorrected. p-cluster values are corrected for multiple comparisons using familywise error (FWE) determined from 100 permutations, p(FWE) < 0.05. Thresholds for cluster extents: 39 and 41, respectively.**

**Table S3. Coupling N2 amplitude frontocentral sensor and BOLD signal**

| | | MNI Coordinates | | | | | | |
|---|---|---|---|---|---|---|---|---|
| **Contrast and Brain Regions** | ~BA | x | y | z | Z Score | T Score | Cluster extent | p-uncorrected |
| **No-Think > 0** | | | | | | | | |
| **Left FG/occipital cortex** | 37 | -36 | -49 | -10 | 4.52 | 5.82 | 359 | < 0.001 |
| **Right SFG/medial PFC/ACC** | 9 | 18 | 56 | 32 | 4.05 | 4.96 | 69 | < 0.001 |
| **Right occipital cortex** | 18 | 36 | -85 | -4 | 3.50 | 4.07 | 136 | < 0.001 |
| **Left SFG/MFG** | 10/46 | -24 | 56 | 17 | 3.43 | 3.97 | 67 | < 0.001 |
| **Right angular gyrus** | 39 | 48 | -61 | 32 | 3.04 | 3.42 | 73 | < 0.001 |
| **No-Think > Think** | | | | | | | | |
| **Right occipital cortex/FG** | 18/19 | 36 | -85 | -7 | 4.15 | 5.14 | 115 | < 0.001 |
| **Right SFG/ACC** | 9 | 15 | 50 | 29 | 3.95 | 4.80 | 227 | < 0.001 |
| **Right/Left PCC/precuneus** | 31 | 3 | -31 | 35 | 3.66 | 4.32 | 209 | < 0.001 |
| **Left MFG/SFG** | 10 | -33 | 59 | 5 | 3.62 | 4.26 | 54 | < 0.001 |
| **Right angular gyrus** | 39 | 42 | -73 | 38 | 3.33 | 3.83 | 104 | < 0.001 |

**BA, Brodmann's area; ACC, anterior cingulate cortex; FG, fusiform gyrus; MFG, middle frontal gyrus; PCC, posterior cingulate cortex; PFC, prefrontal cortex; SFG, superior frontal gyrus. Cluster extents are indicated in number of voxels. Initial p-voxel threshold at $p < 0.005$ uncorrected. p-cluster values are corrected for multiple comparisons using familywise error (FWE) determined from 100 permutations, $p(FWE) < 0.05$. Thresholds for cluster extents: 35 and 40, respectively.**
